# Why Na+ has higher propensity than K+ to condense DNA in a crowded environment

**DOI:** 10.1101/2023.05.15.540899

**Authors:** Egor S. Kolesnikov, Ivan Yu. Gushchin, Peter A. Zhilyaev, Alexey V. Onufriev

## Abstract

Experimentally, in the presence of crowding agent polyethylene glycol (PEG), sodium ions compact double-stranded DNA more readily than potassium ions. Here we have used molecular dynamics simulations and the “ion binding shells model” of DNA condensation to provide an atomic level picture that explains the observed variations in condensation of short (25 base pairs) DNA duplexes in solutions containing different monovalent cations and PEG; several predictions are made. In general, there are two major modes (shells) of ion binding to DNA, internal and external, distinguished by the proximity of bound ions to the helical axis. Externally bound ions contribute the most to the ion-induced aggregation of DNA duplexes. The simulations reveal that for two adjacent DNA duplexes, as well as for a single DNA duplex, the number of externally bound Na^+^ ions is larger than the number of K^+^ ions over a wide range of NaCl and KCl concentrations in the presence of PEG, providing a qualitative explanation for the higher propensity of sodium ions to compact DNA under crowded conditions. The qualitative picture is confirmed by an estimate of the corresponding free energy of DNA aggregation in the presence of different ions: the aggregation free energy is at least 0.2*k_B_T* per base pair more favorable in solution with NaCl than with KCl, at the same ion concentration. The estimated attraction free energy of DNA duplexes in the presence of Na^+^ depends on the DNA sequence noticeably: we predict that AT-rich DNA duplexes are more readily condensed than GC-rich ones in the presence of Na^+^. The sequence dependence of the DNA aggregation propensity is nearly absent for K^+^. Counter-intuitively, the addition of a small amount of crowding agent with high affinity for the specific condensing ion may lead to the weakening of the ion-mediated DNA-DNA attraction, shifting the equilibrium away from the DNA condensed phase.

## 1. Introduction

Condensation of nucleic acids (NAs) has a decisive role in their packaging into cell nucleus,^1^ viruses,^2, 3^ and gene therapy drugs.^4, 5^ In particular, some of the recently developed vaccines against SARS-CoV-2 consist of lipid nanoparticles with condensed mRNA inside.^6–8^ Eukaryotic cells must fit meters of their DNA into the cell nuclei that are only several microns across:^9^ the necessary amount of DNA compaction in the chromatin is achieved via multiple levels of structural organization, starting with the nucleosome^1, 10, 11^ – a complex of positively charged histone proteins with *∼* 150 base pairs of DNA, surrounded by an atmosphere of diverse ions.^12^

Outside of the cell, DNA can also be condensed in solution without the histones, by adding appropriate counterions.^13–26^ In an aqueous solution, DNA condenses only in the presence of multivalent ions with charge +3*e* or higher,^13, 14^ but the addition of a crowding agent such as polyethylene glycol (PEG)^15, 27^ can promote DNA condensation by mono- or divalent cations.

The effect of crowding on the structure and dynamics of different macromolecules has been widely investigated.^27, 28^ Addition of a crowding agent, in particular, facilitates protein folding.^28–31^ Crowding agents non-specifically decrease the solvent volume available for crowded macromolecules. In particular, polyethylene glycol (PEG) is widely used as a crowding agent in experiments related to protein folding and DNA condensation in crowded environment.^15, 27, 32, 33^ According to the model of Ψ-condensation of DNA, PEG and DNA are non-miscible; the addition of PEG causes a decrease of the volume available for the DNA, thus promoting the DNA condensation.^34^

In the crowded environment of a cell nucleus, the state of DNA compaction is determined by a complex interplay^35^ between the histones, some of which are modified, nucleosome repeat length, and various poly- and counterions, including monovalent species such as K^+^ and Na^+^. Understanding the details of this interplay, as well as dissecting it to determine the relative role of each component is critical to the development of a mechanistic picture that connects chromatin structure to its function. Specifically, what is the role of the monovalent ions in determining the shape of chromatin structures? Since it is the balance between electrostatic same charge repulsion and opposite charge attraction that plays the main role here, one might expect that monovalent ions simply provide the non-specific Debye screening, in which case the two ion species would be completely interchangeable. However, the reality is more complex: an intriguing finding was recently reported^36^ that Na^+^ was more efficient than K^+^ in amplifying self-association of nucleosome arrays induced by Mg^2+^ *in vitro*, at concentrations approximating those expected *in vivo*. The same type of non-equivalence between sodium and potassium ions was seen in the highly related context of DNA condensation: Na^+^ was found to be more efficient than K^+^ at condensing DNA fragments under crowded conditions *in vitro*.^15^ A “DNA + counterions” system is much smaller and less complex than nucleosomal arrays, making it more amenable to long enough, fully atomistic Molecular Dynamics (MD) simulations that can provide fine details of the mechanism behind the non-equivalence – details that may otherwise be hard or impossible to infer from lower resolution methods. Our main goal here is to provide these details, leading to a mechanistic picture of DNA condensation by sodium vs. potassium in the presence of an appropriate crowding agent. Specifically, what is the physics behind the difference in the condensation efficiency of sodium vs. potassium seen in the experiment of Zinchenko et al.^15^?

The distinction between sodium and potassium in the context of the DNA, chromatin components, and their multiple functions is potentially highly important, but not always appreciated.^36, 37^ Indeed, the concentration of K^+^ ions is several times higher (up to five fold^38^ according to some estimates) than that of Na^+^ inside the cell nucleus,^38^ while in the extracellular fluids the relative abundance is the opposite, with Na^+^ being the dominant ion.^39, 40^ Differences between Na^+^ and K^+^ interactions with DNA were previously examined using MD simulations^41, 42^ and found to be significant. The distinction between Na^+^ and K^+^ distributions around DNA is caused by 3 factors: smaller Na^+^ ions better fit to DNA binding sites; Na^+^ has smaller radii and approaches closer to DNA surface, which lead to stronger attraction; K^+^ forms clusters with Cl*^−^* more readily, due to a smaller dehydration penalty, which leads to better electrostatic screening of K^+^ charge.^42^ Thus, one can not automatically assume Na^+^ and K^+^ are interchangeable with respect to molecular structures and processes in the nucleus. Yet, aqueous solvents used in relevant experiments and simulations often contain Na^+^ ions^37, 43^ instead of K^+^.

Theoretical and experimental studies of nucleic acids condensation provide a general physical picture of DNA condensation process. The commonly accepted view is that the DNA-DNA attraction is caused by electrostatic interactions.^44–46^ Mean-field theories such as the Debye–Hückel and Poisson-Boltzmann model can not describe the attraction between same charge objects.^13, 21, 47–51^ To describe the nuances of interaction between NA strands, more complex theoretical models were suggested, including highly simplified models that went beyond the mean-field description.^21–23, 49, 52^ Several other models included aspects of the DNA helical geometry.^53–55^ Among theoretical models that examine DNA-ion systems, atomistic Molecular Dynamics (MD) simulations ^56–67^ provide the ultimate level of details that can be afforded within a classical description. These models were used in the past to examine the mechanisms behind DNA condensation.^16, 21, 23, 47, 51, 64, 68–76^ However, atomistic models are still computationally expensive, and arguably not as well suited for analysis of the basic physics of the phenomenon as are simplified, coarse-grained physics models.^77, 78^ Specifically, estimating the potential of mean force^79^ (PMF) between two DNA duplexes^16, 69, 71, 72, 75^ is often very useful, but may become prohibitively expensive for reasonably long duplexes needed to approximate experimental conditions relevant to this work. Instead, we choose to employ a semi-quantitative model briefly described below.

### 1.1. The multi-shell model of ion-induced NA condensation

A general, and relatively simple model (the “multi-shell model”) of NA condensation was developed,^64^ based on a detailed analysis of MD-simulated ionic atmosphere around DNA and RNA duplexes and actual condensation experiments. In this model of ion-induced nucleic acid condensation, the volume around the DNA helical axis is divided into three concentric ion binding shells: one that includes the deeply bound ions (0-7 Å from the helical axis), internal (7-12 Åfrom the axis), and external (12-16 Å) shells,^64^ see Figure 1. It was found that the probability of NA condensation is proportional to the number of ions in the external ion binding shell.^64^ The model explained experimentally observed differences between DNA and RNA condensation.^64^ It was later found consistent with the NA-NA PMF estimates derived directly from atomistic simulations.^16, 69, 71, 72^

**Figure 1:**
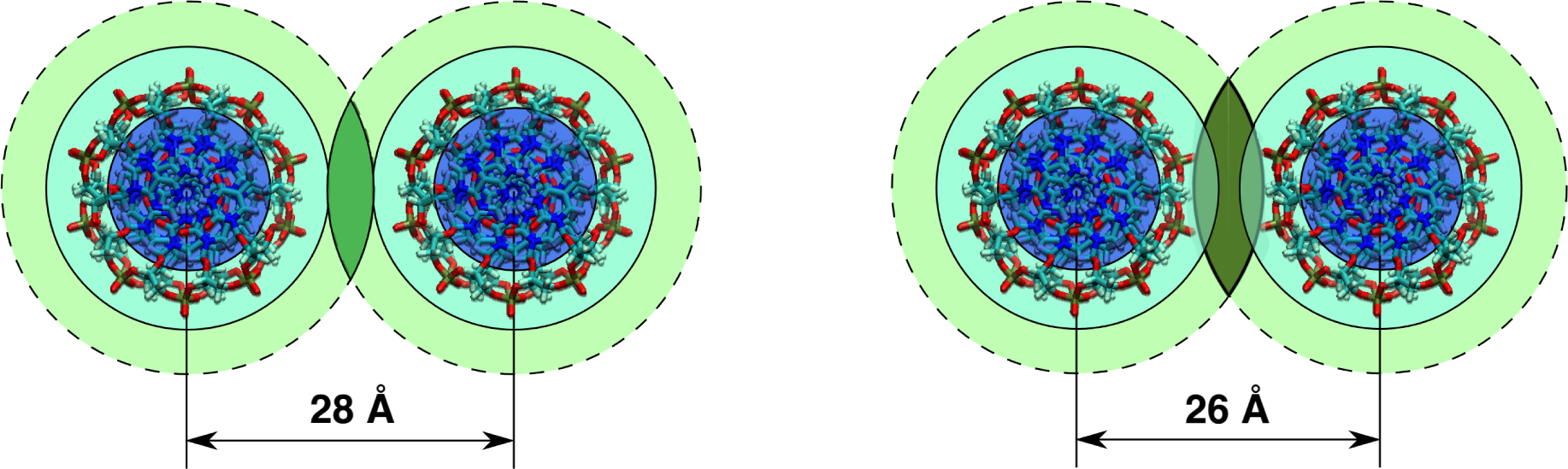
The basis of the multi-shell model of counterion-induced condensation of nucleic acids. Shown are examples of the overlapping ion binding shells of adjacent DNA duplexes corresponding to inter-duplex distances of 28 and 26 Å, respectively. The ions in the dark area cause the attraction between the adjacent duplexes. (a) Overlapping of the external ion binding shells. The dark green area shows the intersection volume between external shells. (b) Overlapping of both internal and external ion binding shells. The darkest green area shows the intersection between the external shells.

In the original multi-shell model, two different regimes of the overlapping of the ion binding shells were distinguished: external-external and internal-external, see Figure 1. Later, the model was supplemented with quantitative expressions that connect the free energy of DNA duplexes attraction in water-ion solution with the number of ions in the intersection of their external ion binding shells.^68^ According to the approach, ion-induced aggregation free energy can be represented as a sum of three additive components:

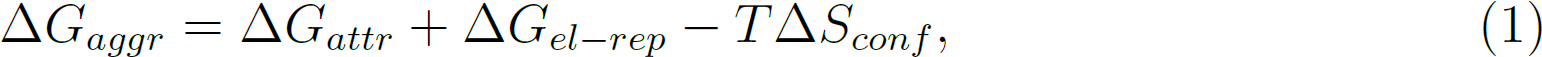

where Δ*G_attr_ <* 0 is a short-range net attractive term, associated with interaction of ions with the oppositely charged DNA duplexes. This attraction is produced by ions accumulated in the intersection of ion binding shells of DNA duplexes. The next term in equation 1 is Δ*G_el−rep_ >* 0, which represents the repulsion between the duplexes, also taking into consideration screening of DNA charges by ions that are not in the intersection of ion binding shells, and the last term, Δ*S_conf_ <* 0, describes the loss of duplex configurational entropy (translational and rotational) due to the aggregation.^68^ While the multi-shell model is a coarse-grained, basic physics model, it has distilled into its framework findings from allatom simulations of counterion-induced NA condensation. The main attractive feature of the model for this study is that it allows one to explore, semi-quantitatively, various aspects of counterion-induced NA aggregation without the need to perform multiple, still very expensive PMF calculations on the DNA duplexes.

Here we use the multi-shell model to understand the physical reason behind the experimentally observed difference between concentrations of NaCl and KCl necessary to induce DNA condensation.^15^ We explore how the numbers of ions in the external ion binding shells of DNA duplexes depend on the ion type: Na^+^ vs. K^+^. We use the model to estimate the DNA-DNA attraction free energy resulting from the accumulation of the cations around DNA duplexes. Based on these estimates we explain the experimental trends and make several testable predictions.

## 2. Results and Discussion

To explain the experimentally observed differences in the concentrations of NaCl and KCl needed for inducing DNA condensation^15^ in the presence of PEG, and to make further predictions, we have investigated the distributions of sodium and potassium ions in the external ion binding shell of DNA duplexes. First, we examined the behavior of sodium and potassium around a single DNA duplex in the presence of 0.5 M NaCl and KCl. Then, we examined the distributions of ions in the intersection of the ion binding shells belonging to two adjacent different duplexes, Figure 1; specifically, we analyzed how the number of ions in the intersection depends on the salt concentration in the range from 0.15 to 2 M. In our simulations, the pairs of duplexes were weakly restrained to structures with minor grooves located opposite to each other; the restraining forces were weak enough to allow the duplexes to move relatively unimpeded within a few Å of the expected inter-duplex distance in the condensed DNA state (24 Å). The initial mutual orientation of the duplexes is similar to the one in the condensed state of the DNA described in Ref.^69^ This orientation also corresponds to the mutual orientation of the DNA grooves in the nucleosome: minor grooves of the turns of the DNA in the nucleosomes are located close to each other and form “supergrooves”.^80^ To simulate experimental conditions of Ref.^15^, 230 g/L of PEG 194 (molecular weight 194 a.m.u.) was added to the solvent, see Methods. All the simulations carried out in this work along with their short identifiers are described in the Supplementary Information, Table S1. The identifier of each simulation has the following structure: 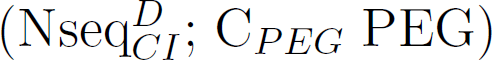, where N is the number of DNA duplexes; seq is the type of DNA sequence; C is the salt concentration; I is the ion type; D is the reference distance for simulations of paired DNA duplexes and C*_PEG_* is the PEG concentration (if C*_PEG_* is missing, it equals to 230 g/L). For example, 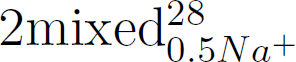 means simulation of 2 mixed sequence DNA duplexes in the presence of 0.5 M Na NaCl and 230 g/L PEG restrained to the structure with inter-duplex distance 28 Å, while (1polyA_0.5_*_K_*^+^; 0 PEG) means simulation of 1 polyA duplex in solution with 0.5 M KCl and no PEG. The relevant characteristics of the computed ion distributions become the inputs into the multi-shell model of DNA condensation, which we use to estimate the change of the free energy of the two-duplex system caused by distribution of Na^+^ and K^+^. These estimates allow us to investigate the dependence the DNA-DNA attraction upon the type of neutralizing ion, to finally address the question of why Na^+^ is more potent than K^+^ at condensing the DNA under crowded conditions. Then we carried out calculation experiment to observe DNA condensation in the presence of monovalent ions and investigated if the numbers of monovalent ions depend on DNA sequence. Finally, we examined the dependence of the numbers of ions in the external ion binding shell on the concentration of the crowding agent to make predictions.

To investigate the difference in the DNA condensation free energy, Δ*G_aggr_*, see equation 1, in the presence of sodium vs. potassium, we estimate the value of ΔΔ*G_aggr_* = Δ*G_aggr_*(*Na*^+^)*−*Δ*G_aggr_*(*K*^+^). Substituting here the definition of Δ*G_aggr_*, see equation 1, we obtain

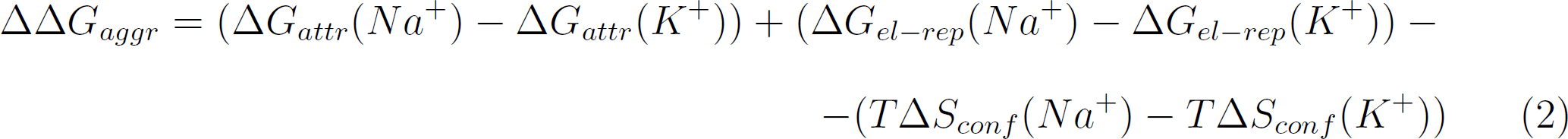

Let’s examine each of the three terms in parentheses in the above expression separately. The DNA-ion systems explored here contain PEG 194; the presence of the crowding agent can affect every term of ΔΔ*G_aggr_*, see equation 2. In particular, PEG is known to interact specifically with proteins, which leads to partial compensation of its crowding effect;^31^ a similar effect may decrease the DNA condensation effect of PEG due to compensation of available volume reduction, thus affecting the *T* Δ*S_conf_* term. Next, according to the model of DNA Ψ-condensation, PEG and DNA are non-miscible, which causes a decrease in available volume for the DNA and leads to DNA condensation.^34^ This effect does not depend on the ion type, so PEG does not affect the last term (*T* Δ*S_conf_* (*Na*^+^) *− T* Δ*S_conf_* (*K*^+^)). In a simulation, the configurational entropy of the DNA duplexes is defined by the conditions of the simulation; in our case, the movement of duplexes was limited by restraints during the simulations. Since the DNA duplexes in different solutions were restrained to the same reference positions with the same restraining harmonic forces with the same constants, Δ*S_conf_* (*Na*^+^) = Δ*S_conf_* (*K*^+^), and thus (*T* Δ*S_conf_* (*Na*^+^) *− T* Δ*S_conf_* (*K*^+^)) = 0. In reality, similar ions are unlikely to affect the configurational entropy of the DNA very differently, so (*T* Δ*S_conf_* (*Na*^+^) *− T* Δ*S_conf_* (*K*^+^)) *≈* 0 is still a reasonable assumption.

Therefore, electrostatic interactions between DNA duplexes and ions, which define Δ*G_attr_* and Δ*G_el−rep_*, contribute the most to the aggregation of DNA duplexes:

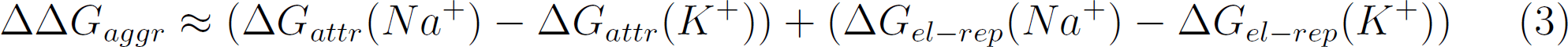

### 2.1. Estimates based on the distribution of ions around a single DNA duplex in the presence of PEG

In this section, we have estimated the propensity of Na^+^ and K^+^ ions to condense DNA by examining their distributions around a single 25 bp long DNA duplex in the presence of PEG. Simulations performed: 1polyA_0.5_*_Na_*+, 1polyA_0.5_*_K_*+, see Table S1 and Methods. The number of sodium ions in the external ion binding shell of DNA calculated from simulations (28.4 *±* 0.2) is significantly higher than that of potassium (22.3 *±* 0.2).

To complete the estimation of ΔΔ*G_aggr_*, we continue examining equation 3. First, we consider Δ*G_el−rep_*(*Na*^+^) *−* Δ*G_el−rep_*(*K*^+^) in the presence of crowding agent we use. PEG is a neutral crowding agent, and its contact with DNA is thermodynamically unfavorable. Therefore, PEG, not being in proximity of attracting DNA duplexes, can not affect Δ*G_el−rep_*(*Na*^+^)*−*Δ*G_el−rep_*(*K*^+^). In the simulations with a single DNA duplex, Δ*G_el−rep_* represents hypothetical repulsion energy between two DNA duplexes approached to each other. It can be affected by the degree of DNA charge neutralization by ions, that do not contribute to Δ*G_attr_* of adjacent DNA duplexes. In our simulations, the degree in the presence of sodium (86%) is higher than that of potassium (80%), thus Δ*G_el−rep_*(*Na*^+^) *−* Δ*G_el−rep_*(*K*^+^) *<* 0. Therefore,

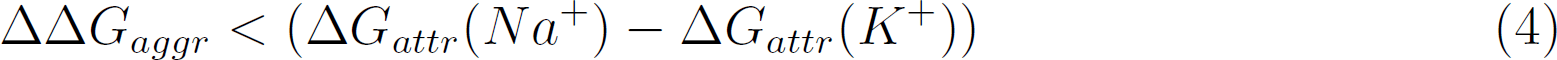

The attraction term can be estimated via the multi-shell model (see Methods) using the numbers of cations in the external ion binding shell of DNA from the simulations as an input. If all the ions in the external shell contribute to Δ*G_attr_*, in the system with sodium chloride, Δ*G_aggr_* = *−* 33.2 *±* 0.3*k_B_T*, which is less than that in the system with potassium, Δ*G_aggr_* = *−*20.5 *±* 0.2*k_B_T*. These values represent the aggregation energy per 25 bp long polyA duplex, assuming that all the ions in the external ion binding shell contribute to Δ*G_aggr_*. To make connection to reality, one has to account for the realistic arrangement of duplexes in the condensed state, and the sharing of the ions between the ovelapping external binding shells, see Figure 1a. Experimentally, DNA is found to condense into hexagonally packed structures in the presence of multivalent ions. ^81^ So, in the intersection, only *≈* 1*/*6 of externally bound ions contribute to attraction from both of the duplexes. Thus ΔΔ*G_aggr_ <* (Δ*G_attr_*(*Na*^+^) *−* Δ*G_attr_*(*K*^+^))/3 = *−*4.2 *±* 0.1*k_B_T*, where the factor of 3 accounts for half of the attractive interactions with the six nearest neighbors in the hexagonally packed aggregate. This value is equal to *−*(0.17 *±* 0.01)*k_B_T* per base pair, which is in the range expected from experiment.^82^ The difference between the aggregation energy of the DNA duplexes indicates that at a certain salt concentration the aggregation energy is enough for the DNA condensation to begin in the presence of sodium, but not in the presence of potassium. The free energy of the system depends strongly on the ion type, with Na^+^ being more potent than K^+^ at condensing the DNA. Thus, we have established that the attraction between two DNA duplexes depends on the type of ion present in the solution. This phenomenon can be analyzed by employing multi-shell model of NA condensation. Different ions have different abilities to shield the negatively charged phosphate backbone of the DNA and promote DNA-DNA attraction.

#### 2.1.1. Qualitative validation of the model: the effect of Rb**^+^**

In the experiment by Zinchenko^15^ et al., DNA condensation in the presence of PEG was also induced by RbCl. The experiment showed that Rb^+^ was less potent to condense DNA than K^+^. This observation provides an independent test of our theory: can we predict the correct relative propensity of Rb^+^ to condense the DNA in a crowded environment? To this end, we have repeated the single-duplex simulation of polyA DNA described above with RbCl. Simulations performed: 1polyA_0.5_*_Na_*+, 1polyA_0.5_*_K_*+, and 1polyA_0.5_*_Rb_*+. The number of Rb^+^ ions in the external ion binding shell of DNA is found to be less than that of K^+^, see Table 1; according to the multi-shell model, the propensity of ion to condense the DNA is proportional to the number of ions in the external ion binding shell. This comparison explains the difference of K^+^ and Rb^+^ propensity to condense DNA.

**Table 1:** Numbers of different ions in the external ion binding shell of polyA DNA in the presence of PEG correlate with probability of DNA condensation in the experiment.^15^ According to the model used in this work, a larger number of ions corresponds to higher propensity to condense DNA. The ions are listed in the order of decreasing experimental propensity to condense DNA in the presence of PEG;^15^ the predicted ordering based on the numbers of ions in the external shell (second column) agrees with experiment. Bulk salt concentration is 0.5 M.

| Cation type | Number of ions in the external ion binding shell |
| --- | --- |
| Na<sup>+</sup> | $28.4 \pm 0.2$ |
| K<sup>+</sup> | $22.3 \pm 0.2$ |
| Rb<sup>+</sup> | $20.8 \pm 0.2$ |

Note that a qualitative validation such as the one presented above is sufficient for the purposes of this work that aims to provide qualitative explanations and predict qualitative trends.

### 2.2. Ion distributions between two adjacent DNA duplexes and their attraction

In the previous section, we estimated the propensity of ions to condense DNA based on distributions of ions obtained from simulations of a single DNA duplex, which allowed us to make qualitative conclusions. The approach was justified previously^68^ for multivalent ions, which bind strongly to the DNA. However, monovalent ions are relatively mobile, so they may redistribute between two adjacent DNA duplexes as these come close to each other. To investigate this possibility, we have explored ion distributions between two DNA adjacent duplexes. Here, we examined a pair of 25 bp polyA DNA duplexes submerged in a rectangular simulation box of the appropriate solvent, see Methods. We hypothesized that the phosphate groups of adjacent duplexes may form the cation-bridged inter-DNA contacts^69^ differently in the presence of sodium vs. potassium. Previously, the condensation of the effectively infinite DNA fibers was simulated in the presence of NaCl and MgCl.^69^ The observed inter-duplex distance in the condensed state was 24 Å, and it did not depend on the ion type (Na^+^ vs. Mg^2+^). Accordingly, here we have assumed that in the presence of sodium and potassium the inter-duplex distances in condensed state do not differ noticeably. Based on this assumption, we simulated the pairs of duplexes with inter-duplex distance constrained to 24 Å. The initial mutual orientation of duplexes is the same as in Ref.,^69^ with the minor grooves of adjacent DNA duplexes located opposite to each other. We have performed a series of MD simulations, in which the DNA duplexes were harmonically restrained strongly enough to prevent the minor grooves of adjacent DNA duplexes from moving away from each other. But, at the same time, the restraining forces were chosen to be weak enough to allow the inter-duplex distance to fluctuate easily between 22 and *∼* 28 Å. Consequently, we were able to determine the values substantial for our calculations: differences in the number of ions between DNA duplexes, and fluctuations of the inter-duplex distance. To reach the middle point of the DNA condensation transition, we have simulated solutions with high concentrations of sodium and potassium chlorides (0.5, 1 and 2 M) and with physiological concentration (0.15 M) in different systems. Here we have used higher monovalent salt concentrations than used in the experiments of Ref.,^15^ because the experiments showed that the shorter the crowding molecules are, the bigger the minimum salt concentration is needed to condense DNA. So, since we used low-molecular weight PEG in our simulation, higher salt concentrations are needed to induce DNA condensation. Simulations performed: 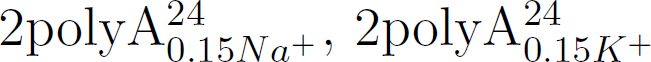, 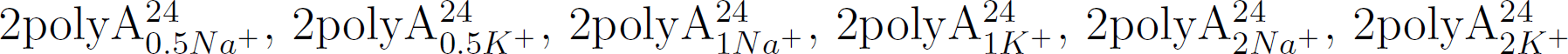,

#### 2.2.1. Densities and numbers of Na^+^ and K^+^ ions between adjacent DNA duplexes

The main qualitative outcome of the simulations described above is shown in Figure 2: more sodium ions accumulate between adjacent DNA duplexes than potassium ions, and this difference leads to a stronger DNA-DNA attraction in the presence of sodium compared to potassium. Thus, the qualitative picture developed above, which was based on computed ion distributions around a single duplex, remains valid.

**Figure 2:**
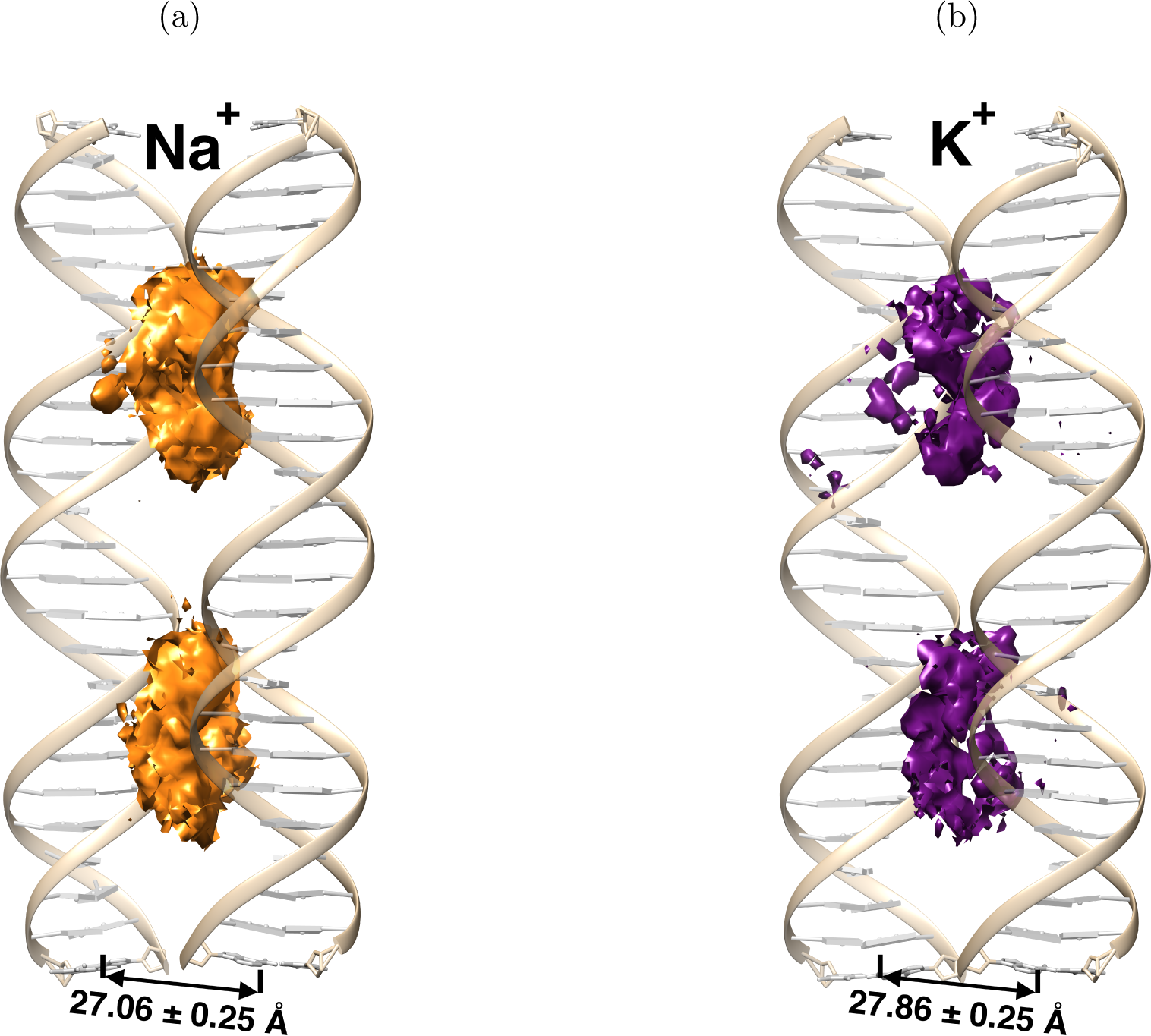
The computed densities of sodium (orange, left) and potassium (purple, right) ions between two adjacent DNA duplexes in solutions with 0.15 M salt concentration. Shown are the parts of maps, at the level of 3.5 electron charges per Å^3^, which are no further than 16 Å away from both of the helical axes. The concentration of Na^+^ ions between adjacent DNA duplexes is noticeably higher than that of K^+^. During the simulations, the duplexes were weakly restrained to reference structures, with the inter-duplex distance of 24 Å. The equilibrium distance between the duplexes during simulations for each ion type is shown under the maps, it does not depend on the ion type significantly within the error bar. Ions tend to concentrate between the duplexes near their minor grooves. Sodium accumulates better than potassium between adjacent DNA duplexes.

Monovalent cations reside in the negatively charged pockets formed by phosphate groups of adjacent duplexes. These results are consistent with previous findings.^69^ Sodium accumulates between DNA duplexes better than potassium.

To quantify the difference in the numbers of sodium and potassium ions accumulated between DNA duplexes, we counted the number of ions in the intersection of the ion binding shells (Figure 1a). These ions interact with both of the duplexes and cause their attraction. The calculated numbers of ions in the intersection of the shells are presented in Figure 3 and Table S2.

**Figure 3:**
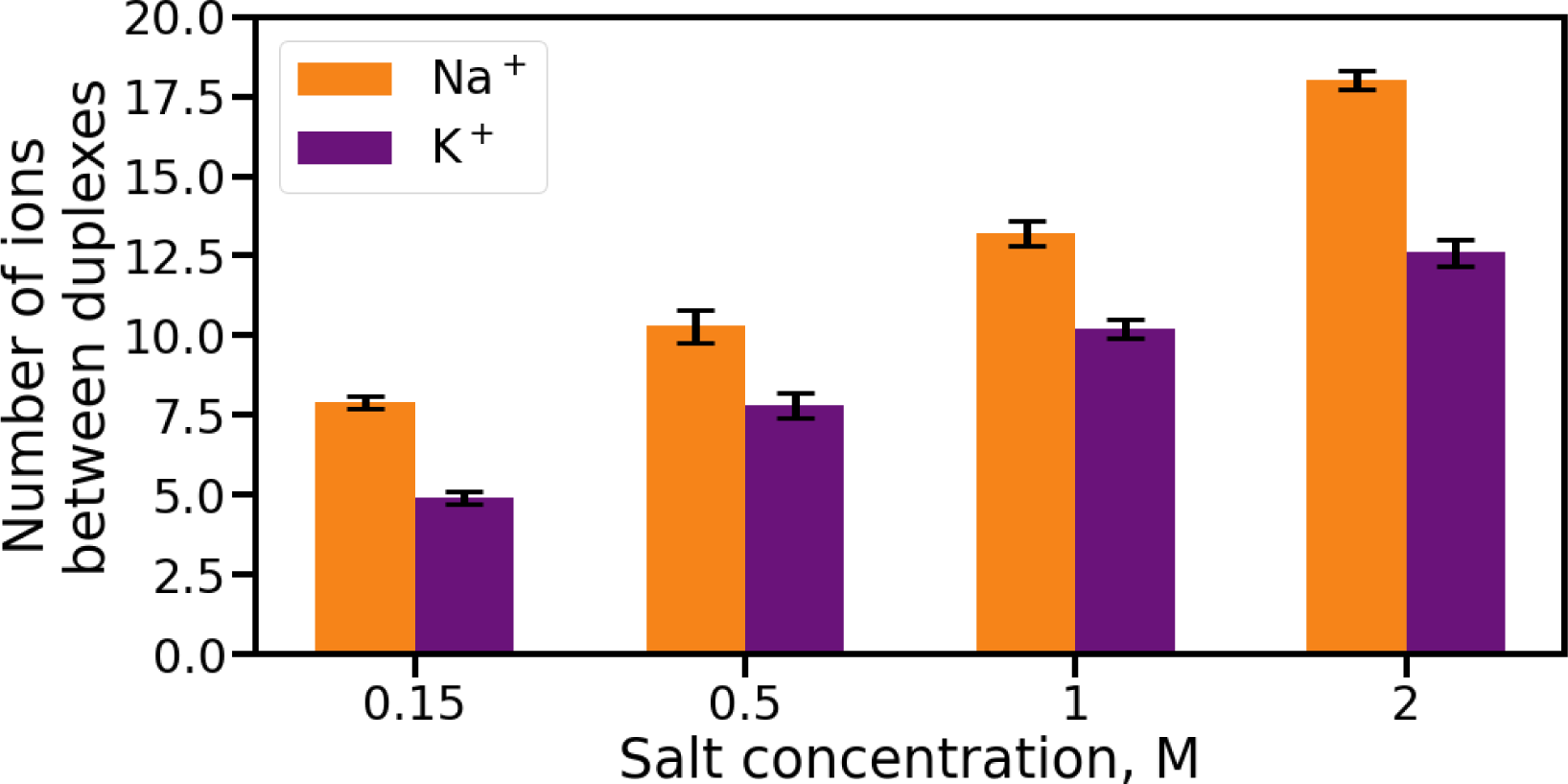
The number of sodium ions in-between two adjacent DNA duplexes is always higher than that of potassium, in a wide range of salt concentrations. Shown are the numbers of Na^+^ and K^+^ ions in the intersection of the ion binding shells (Figure 1) of two adjacent DNA duplexes. Error bars indicate the standard errors of the mean, see Methods.

The number of sodium ions between DNA duplexes is higher by about 30-60 % than that of potassium for all of the concentrations tested. The estimated numbers of ions in the intersections of the external ion binding shells with theinternal shells is presented in Table S2. The number of sodium ions taking part in internal-external interaction is higher than that of potassium, but they do not differ as much as the whole number of ions in the intersection of the ion binding shells. In summary, the sodium ions have a higher concentration in external-external and internal-external intersections compared to the potassium ions. Both of the intersection regimes (external-external and internal-external) contribute to the DNA-DNA attraction, leading to aggregation of the DNA duplexes. Thus, sodium ions produce stronger attraction in both of the regimes.

#### 2.2.2. Estimation of DNA aggregation energy based on ion distributions around two adjacent DNA duplexes

Here we have again used the multi-shell model of ion-induced DNA condensation to estimate the free energy of aggregation of DNA.^64, 68^ As we discussed earlier, the ions in the intersection accumulate between the phosphate groups of adjacent DNA duplexes (Figure 2). These ions interact with both of the duplexes and produce attraction.^69^ We use equation 3 to estimate ΔΔ*G_aggr_* based on the numbers of ions in the intersection of external ion binding shells of adjacent DNA duplexes. Let’s begin with the repulsive term Δ*G_el−rep_*(*Na*^+^)*−*Δ*G_el−rep_*(*K*^+^). We have calculated only the energy of duplexes’ repulsion and did not take into account charge screening by ions. This difference is less than 0.4 *· k_B_T* in every trajectory, see Table S3. The difference is caused by unequal distances between duplexes in the presence of sodium and potassium. The number of charge screening ions, that are not in the intersection of ion binding shells, in solutions with sodium and potassium do not differ significantly in our simulations, so 0.4 *· k_B_T* is a reasonable upper estimate for Δ*G_el−rep_*(*Na*^+^) *−* Δ*G_el−rep_*(*K*^+^). Thus,

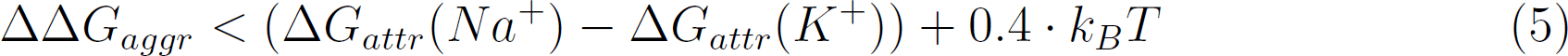

To estimate the DNA-DNA attraction term, Δ*G_attr_* in equation 5, we used the computed numbers of cations in the intersection of ion binding shells, see Figure 3 and Table S2 and Methods. The result, for a range of ion concentrations, is summarized in Figure 4 and Table S4.

**Figure 4:**
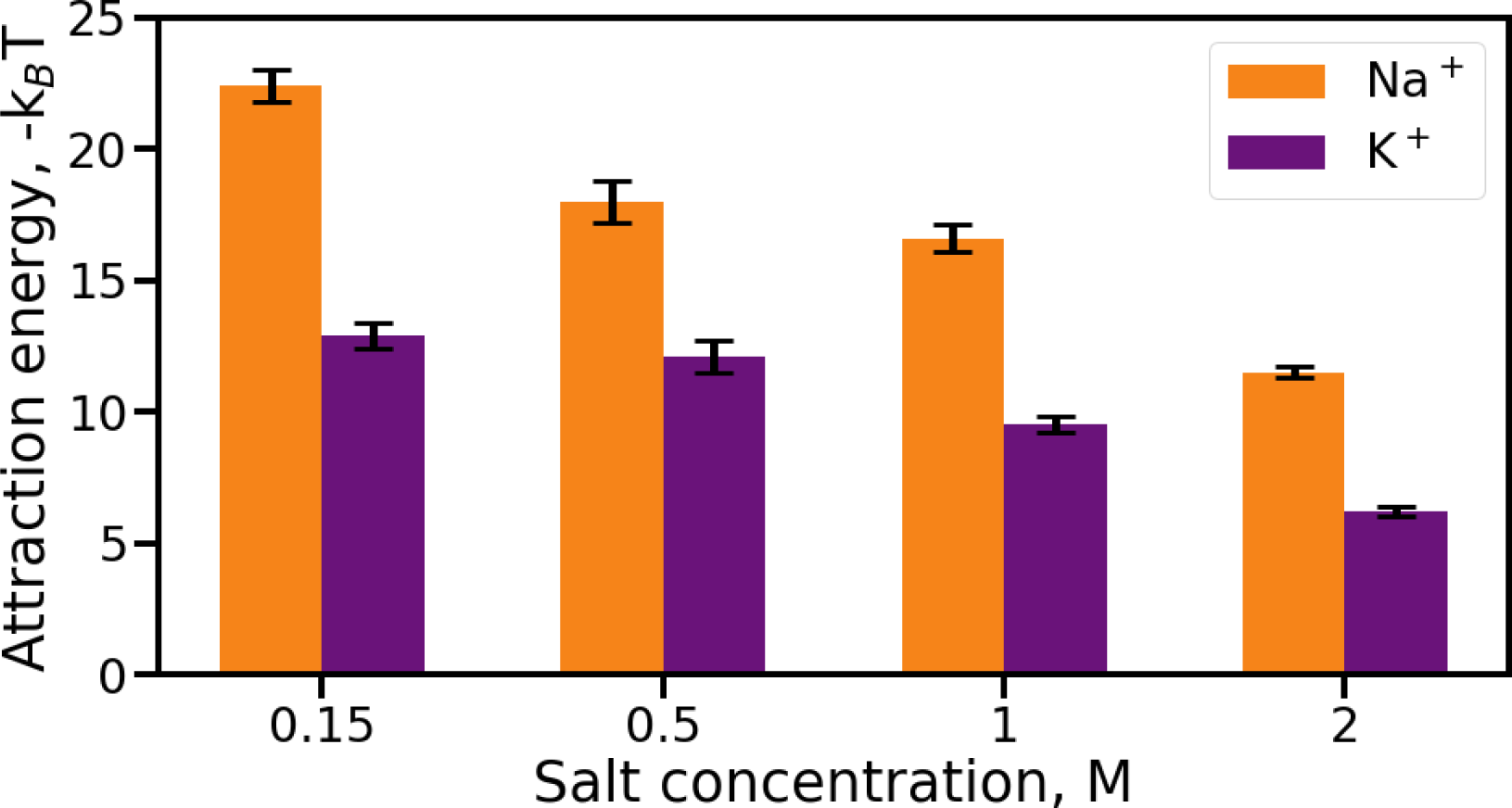
The estimated attraction free energy between two DNA duplexes caused by counterions. In the presence of sodium ions, the ion-induced DNA-DNA attraction is stronger than in solution with potassium, over a wide range of salt concentrations.

In the systems with sodium chloride, Δ*G_attr_* is also less than Δ*G_attr_* in the systems with potassium by at least 5.3 *· k_B_T*. Thus ΔΔ*G_aggr_ < −*(4.9 *±* 0.3) *· k_B_T* in a wide range of salt concentrations. The difference of aggregation free energies per base pair is equal to *−*(0.20 *±* 0.01) *· k_B_T*. The estimation is similar to the one obtained from simulations of a single DNA duplex and it is large enough to induce DNA condensation in the presence of sodium, but not in a solution with potassium with certain concentration of NaCl and KCl. Aggregation energy depends strongly on the ion type, with Na^+^ being more potent in DNA condensation than K^+^. These calculations agree with, and explain the physical origins of the previous experimental results.^15^

A word is due here on the specific choice of the method employed here to estimate the inter-duplex aggregation free energy. A more standard approach, compared to the one used above, could be based on computing the corresponding potential of mean force (PMF).^79^ Various approaches to obtain the PMF are suitable for different systems,^83^ including conformational changes in proteins,^84^ or biophysical events in the active sites of RNA.^85^ The PMF for two DNA duplexes approaching each other was previously calculated under various conditions.^69, 71, 75^ The inter-duplex distance in the condensed state varied from 24 Å to 28 Å in previous studies, depending on the conditions of the simulations. In these estimates, the DNA duplexes were assumed parallel to each other: that assumption would have to be relaxed if one were to fully account for the loss of conformational entropy upon DNA condensation, to obtain the actual aggregation free energy. An averaging over all possible conformational states would have to be performed, which could make the computation prohibitively expensive for long enough duplexes. In the specific case of counterion induced NA condensation, the multi-shell model employed here provides a relatively inexpensive alternative to computing the PMF; the multi-shell model includes an estimate of the loss of conformational entropy upon DNA aggregation, thus yielding an approximation for the total free energy of the DNA aggregation.

#### 2.2.3. Direct observation of attraction between two DNA duplexes in simulation

To further validate our findings based on the estimates of the DNA aggregation free energy, we have simulated two adjacent DNA duplexes, initially separated by 28 Å inter-duplex distance. The duplexes were parallel, mutually oriented as in Ref.^69^ The main idea behind this simulation is as follows. The inter-duplex distance in the condensed state in the presence of NaCl is expected to decrease to 24 Å.^69^ If DNA duplexes attract in the presence of Na^+^ or K^+^, then we expect the average inter-duplex distance during simulation to be less than 28 Å, which duplexes are constrained to. If, on the other hand, the ion atmosphere does not produce strong enough DNA-DNA attraction, then we expect the average distance to remain *≥* 28 Å in the course of the simulation. Simulations performed: 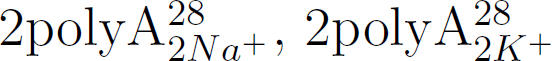.

The trajectory-averaged distances between DNA axes are NaCl: 26.4 *±* 0.4 Å, KCl: 28.16 *±* 0.23 Å. The violin plot (Figure 5) further illustrates inter-duplex distances during the simulations in solutions with NaCl and KCl.

**Figure 5:**
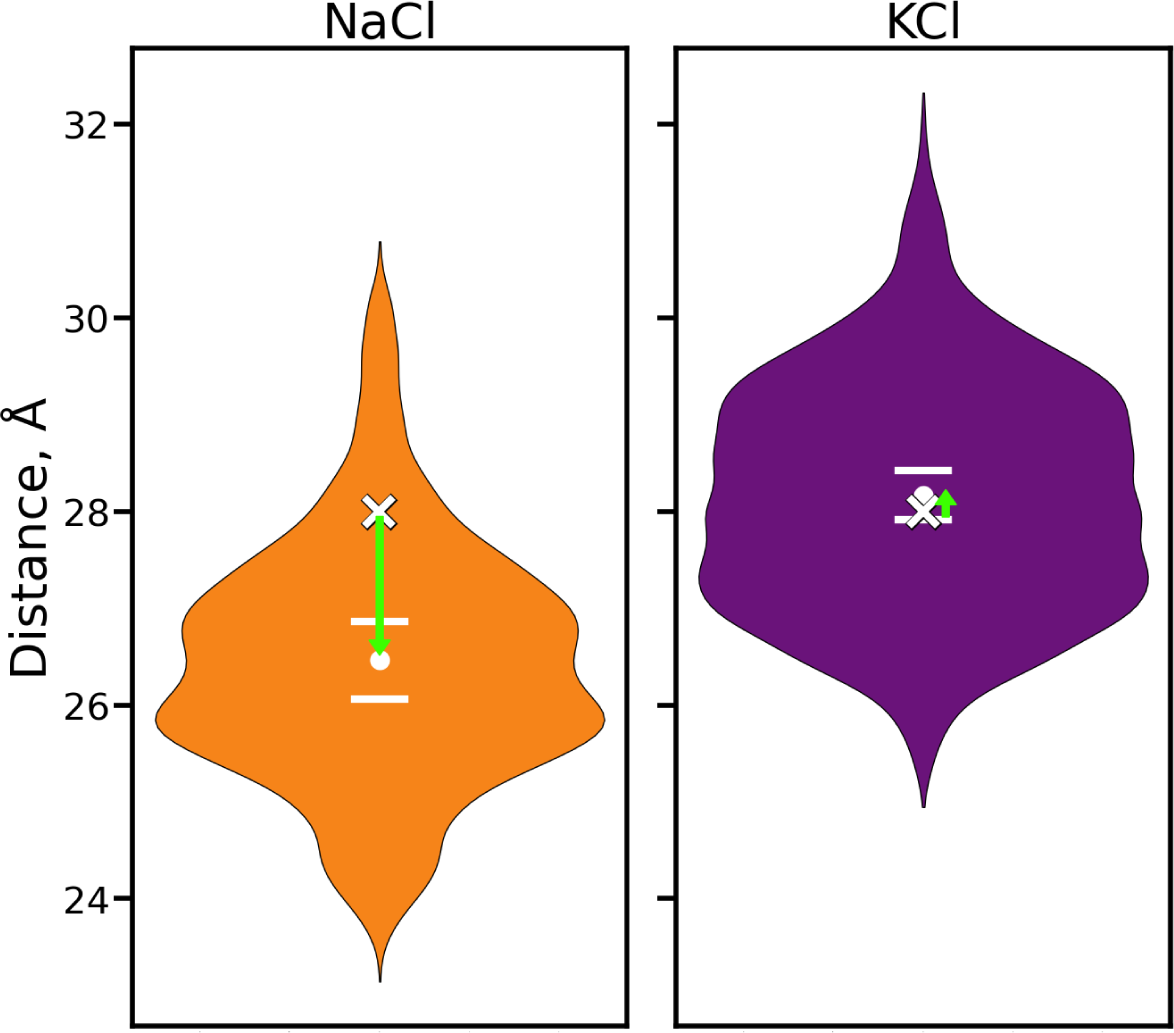
DNA-DNA attraction directly observed in atomistic simulations. Shown are the computed distributions of inter-duplex distances during simulations of DNA duplexes in the presence of NaCl (orange, left) and KCl (purple, right). The initial inter-duplex distance is 28 Å, indicated by the white crosses. Mean distances observed during the simulations are shown as white dots, and standard deviations of the means as white lines. Green arrows show the displacement of the DNA during the simulation: in NaCl the duplexes approach each other, while in KCl the inter-duplex distance remains virtually unchanged. The bulk salt concentration is 2 M.

It can be seen from the plot and the mean values that in solution with NaCl the average inter-duplex distance is appreciably less than 28 Å, but in the presence of KCl it is *≥* 28 Å. This observation indicates that at the same concentration of NaCl or KCl in solution, sodium ions induce DNA condensation (as approximated by the two-duplex system), but potassium ions do not.

### 2.3. Effect of DNA sequence on the condensation

The dependence of DNA condensation on the DNA sequence in the presence of multivalent ions was investigated previously, see *e.g.* Refs.^64, 76^ In particular, AT-rich sequences were found to be condensed more easily than GC-rich ones by a variety of polyions, including cobalt hexamine, spermine and polylysine. It was also previously reported that the distributions of deeply bound Na^+^ and K^+^ ions around double-stranded DNA^41^ depend on the DNA sequence. However, to the best of our knowledge, sequence dependence of DNA condensation by monovalent ions in the presence of crowding agents was not previously examined. To determine if the attraction between DNA duplexes in the presence of monovalent ions may depend on the DNA sequence, here we have examined a pair of 25 bp mixed-sequence DNA duplexes (GCATCTGGGCTATAAAAGGGCGTCG) in different mutual configurations, to compare with polyA DNA explored above.

We have repeated some of the computational experiments described in Sec. 2.2 and 2.3 with this pair of DNA duplexes, weakly restrained to their reference structures separated by the inter-duplex distances of 24 or 28 Å, at bulk salt concentration of 2 M. Simulations performed: 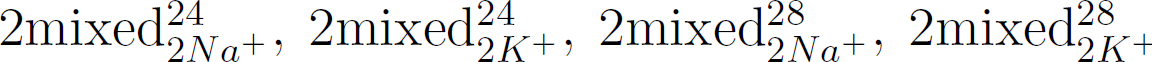 The results, along with the corresponding numbers for polyA DNA, are summarized in Figure 6 and in Tables S7 and S8.

**Figure 6:**
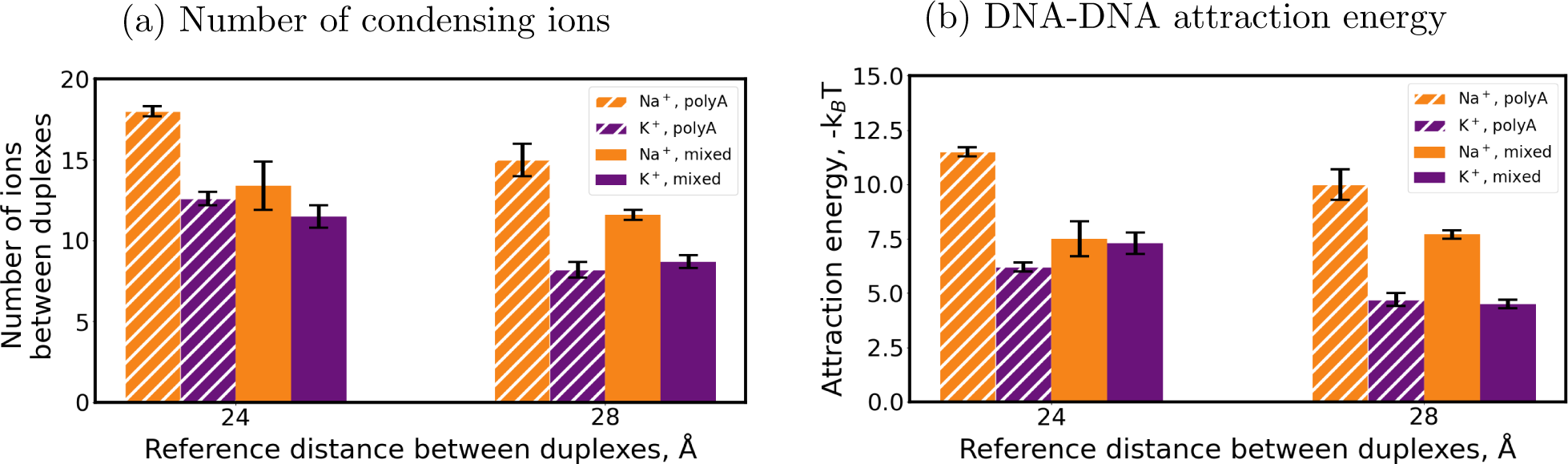
Number of condensing ions and ion-induced DNA-DNA attraction free energy depend strongly on the DNA sequence for Na^+^ ions (orange), but not for K^+^ ions (purple). (a) The number of ions in the intersection of the ion binding shells responsible for DNA condensation, for the mixed sequence (solid bars) and polyA (cross-hatched bars) duplexes. (b) The corresponding inter-duplex attraction free energy. The bulk salt concentration is 2 M.

One can draw three main conclusions from Figure 6 and the supporting information:

First, the number of condensing ions and the ion-induced DNA-DNA attraction free energy depend strongly on the DNA sequence for Na^+^ ions (orange bars), but not for K^+^ ions (purple bars). Consequently, the absolute value of the predicted Δ*G_attr_* decreases noticeably for Na^+^ ions, but not for K^+^, as seen in the Figure 6 in going from the polyA to the mixed sequence duplexes. We therefore predict that the aggregation propensity of DNA in the presence of PEG will depend on the sequence if the condensing counterion is Na^+^ but not K^+^. The general conclusion on the dependence of the number of ions on the DNA sequence is further confirmed by a computational experiment with a single DNA duplex, described in Sec. 2.1, see details and caveats in Supplementary Information.

Second, the observed dependencies of the numbers of Na^+^ and K^+^ ions on the DNA sequence lead to a decrease in the difference between numbers of Na^+^ and K^+^ contributing to condensation when the DNA sequence is switching from polyA to the mixed one. Thus the difference between concentrations of NaCl and KCl inducing mixed DNA condensation is less than that for polyA DNA. This finding is consistent with previous results for condensation in the presence of multivalent ions.^76^ The higher propensity of Na^+^ vs. K^+^ to condense the DNA remains as the DNA sequence is changed. However, the difference in the propensity to condense decreases: in terms of ΔΔ*G*, the difference is decreased *≥* 1.6 fold with 24 Å reference distance.

Third, the sequence dependence trends discussed above are robust to the inter-duplex distance used in the simulations: the same trends seen for each ion type are the same when either 24 Å or 28 Å inter-duplex distance is used in the simulations. Specifically, the number of Na^+^ ions in the intersection of external shells, and the absolute value of attraction energy in the presence of Na^+^, are higher for polyA sequence than for the mixed one in the simulations with both reference distances. We did not see a noticeable difference between K^+^ numbers in the intersection of ion binding shells in the simulations with both reference inter-duplex distances for pairs of duplexes with polyA and mixed sequences.

While a detailed investigation into possible origins of the sequence dependence of DNA condensation by monovalent ions under crowded conditions is beyond the scope of this work, we offer the following insights. The dependence of monovalent ion distributions on the GCcontent of DNA was examined earlier:^41^ a significant fraction of monovalent cations was found to be bound deeply, to specific binding sites near the DNA axis. The occupancies of the specific binding sites near guanine and cytosine were higher than those of binding sites in AT-tracts. Since the total numbers of monovalent ions bound to the DNA is expected to be largely independent of the sequence – see *e.g.* Ref.^41^ for an argument based on counterion condensation theory – it follows that the number of ions in the external shell, most important for condensation, should also be sequence-dependent. Another relevant question is why the sequence dependence of DNA condensation propensity is different between Na^+^ and K^+^? Earlier studies provided a comparative analysis of Na^+^ and K^+^ distributions around mixed DNA^42, 86^ and showed that, compared to potassium, a larger fraction of sodium ions accumulate in the DNA interior.^42^ This distinction is partially caused by a relatively better fit of Na^+^ to the binding sites. The difference between Na^+^ and K^+^ affinities for the deeply buried binding sites is expected to translate into a difference between relative occupancies of the extermnal binding shell, and, hence, into a difference in the condensation propensities. We stress that the above discussion does not provide a definitive “proof”; rather, we have outlined one possible mechansim.

### 2.4. The addition of PEG may have the opposite effect on DNA condensing potency of different ions

The effect of the addition of PEG to solution with DNA is well established experimentally and theoretically.^27, 34^ According to the model of DNA Ψ-condensation, PEG and DNA are non-miscible, which causes the decrease in available volume for DNA and leads to DNA condensation.^34^ Likewise, the addition of PEG decreases the volume available for the ions, also favoring condensation. But note that some ions may bind to the added PEG molecules – these ions will become “sequestered”, will no longer interact with the DNA, and so no longer participate in the ion-mediated electrostatic attraction between DNA molecules, which plays the key role.^55, 64, 68, 69^ In this scenario, the addition of a small amount of PEG will not decrease the volume available for the DNA (and ions) dramatically, but the PEG molecules may still bind enough ions to decrease the ion-mediated attraction force between the DNA duplexes noticeably. The competition between contributions to the DNA attractive free energy from the non-specific crowding, which favor condensation, and from the capturing of ions by the crowding molecules, will determine the outcome. If the affinity of the ions to the crowding agent is high enough, the contribution of the specific ion binding to the crowding agent, which effectively opposes DNA condensation, may be higher than the contribution of non-specific crowding, which favors it. So, the general effect of the addition of PEG, in this case, may be opposite of the expected – DNA will condense less easily.

In what follows we will demonstrate how the above idea may work, quantitatively, for the case of DNA condensation induced by sodium or potassium in the presence of PEG. According to the multi-shell model of ion-induced DNA condensation, the propensity of an ion species to condense DNA is proportional to the number of ions in the external ion binding shell of the DNA duplex. As discussed above, the number of ions in the external ion binding shell may depend on the type of crowding agent added to the solution. This dependence is determined by the affinity of an ion to the crowding agent that modulates the chemical potential of the ion in the solution.

To check if PEG may reduce the numbers of Na^+^ or K^+^ ions in the external ion binding shell of the DNA, we have examined systems containing a DNA duplex with ions in the presence of 28 g/L of PEG, and with no PEG, in addition to the simulations with “default” 230 g/L PEG. Simulations performed: (1polyA_0.5_*_K_*+; 0 PEG), (1polyA_0.5_*_K_*+; 28 PEG), 1polyA_0.5_*_K_*+, (1polyA_0.5_*_Na_*+; 0 PEG), (1polyA_0.5_*_Na_*+; 28 PEG), 1polyA_0.5_*_Na_*+. Numbers of ions in the external ion binding shell of polyA DNA in the presence of different concentrations of PEG are presented below, Figure 7.

**Figure 7:**
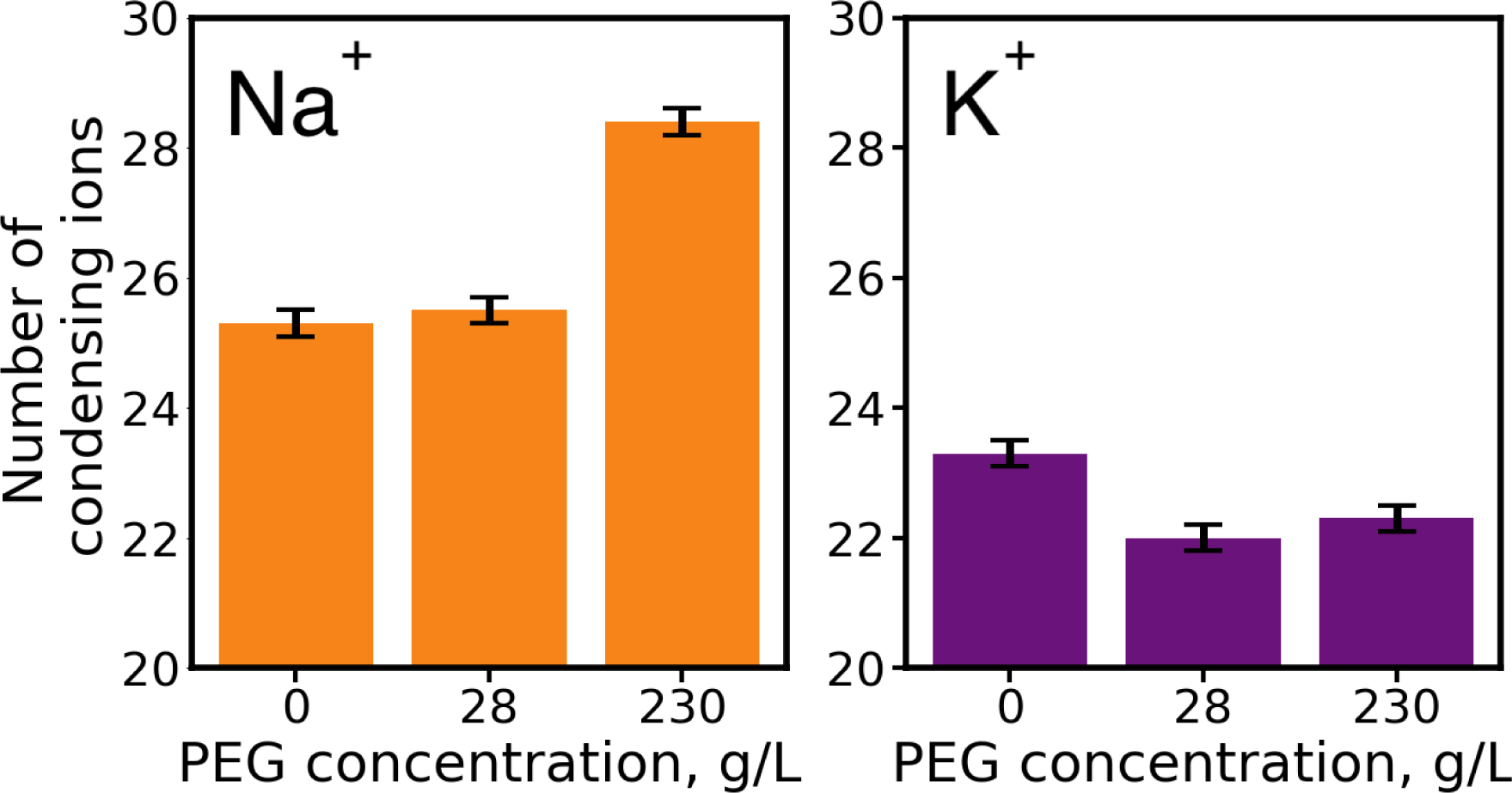
The effect of PEG on the number of ions in the external ion binding shell of the DNA depends significantly on the ion type. Shown are the numbers of ions in the external ion binding shell of polyA DNA in the presence of different PEG concentrations. The number of Na^+^ ions in the external ion binding shell of the DNA increases (left panel) as the PEG concentration increases, but the trend is the opposite for K^+^ (right panel). Bulk salt concentration is 0.5 M.

As one can see from the graph, the number of Na^+^ in the external ion binding shell of DNA (the ions that contribute the most to DNA condensation) increases with growing PEG concentration. This dependence indicates that non-specific crowding of sodium ions from the solution overcomes specific binding of PEG molecules to the ion. In contrast, the number of K^+^ ions slightly decreases with the addition of PEG. This decrease is caused by strong specific binding of potassium ions to PEG that exceeds crowding. So, the numbers of condensing Na^+^ and K^+^ ions depend on the PEG concentration in the opposite way.

The above argument shows that the addition of a small amount of crowding agent to KCl solution with DNA may, counterintuitively, weaken the ion-induced DNA-DNA attraction by decreasing the absolute value of the attractive component of the condensation free energy. It is not obvious to us if this effect can be directly observed experimentally, since monovalent ions condense the DNA only in the presence of PEG at high concentrations. However, a similar effect may be possible to observe directly for divalent ion-induced DNA condensation: DNA condenses in the presence of Mg^2+^ at a low concentration of crowding agent. Likewise, the addition of a crowding agent with high affinity to Mg^2+^ may lead to decondensation of the DNA. Trivalent ions condense DNA by themselves.^81^ So, if the affinity of a multivalent ion to the crowding agent is high enough, the addition of a small amount of the crowding agent may lead to a weakening of the DNA-DNA attraction.

We propose the following experiment to test the above prediction for monovalent ions, albeit indirectly. The addition of PEG to the DNA solution leads to a rise in DNA condensation probability due to the decrease of the volume available to the DNA fibers. This phenomenon does not depend on the ion type. When this effect is combined with ion-induced DNA-DNA attraction, DNA condensation may occur. As one can see from Figure 7, the addition of PEG increase the number of Na^+^ ions that participate in condensing the DNA, but does not increase the number of K^+^ ions causing the DNA condensation. Based on this result, we predict that if one gradually adds PEG to DNA-containing solutions with the same concentrations of NaCl and KCl, at some critical PEG concentration the DNA condensation will start in the presence of sodium, but not potassium.

We conclude this section by a speculation, based on an extension of our reasoning to mutivalent ions. DNA duplexes readily condense with the addition of multivalent ions. The addition of a crowding agent to the solution should decrease attraction energy of DNA fibers, which is negative, and lead to stronger attraction of duplexes. As one can see in Figure 7, for some ions with high affinity to the crowding agent it may be energetically beneficial to unbind ion from DNA when a crowding agent is added. A low enough concentration of crowding agent can cause only a small, non-specific increase in the condensation probability. But the same crowding agent may bind enough ions to reduce their potency to condense DNA significantly. Based on these consideration, we speculate that in an experiment in which just enough multivalent ions were added to cause DNA condensation, subsequent addition of a small amount of crowding agent that has high affinity for the multivalent ion may cause DNA de-condensation.

## 3. Conclusions

In this work we have investigated, theoretically and computationally, various aspects of DNA condensation by monovalent ions in the presence of a crowding agent polyethylene glycol (PEG). Our main tools were atomistic simulations and the multi-shell model of ion-induced DNA condensation, which we employed to estimate free energy components of counterion-induced DNA-DNA aggregation.

Our first goal was to explain earlier experimental results:^15^ sodium ions condense DNA more readily than potassium in the presence of PEG. Our simulations have revealed noticeable differences in the ability of sodium and potassium to accumulate between two adjacent DNA duplexes: the number of sodium ions between adjacent DNA duplexes is 30-60% higher than that of potassium. Thus, more sodium ions accumulate between phosphate groups of adjacent DNA duplexes as compared to potassium ions. Counterions in this location are known to be key to DNA condensation,^64, 69^ which explains, qualitatively, why sodium ions produce stronger attraction of DNA duplexes compared to potassium.

We have made the above argument quantitative by employing the multi-shell model^68^ of ion-induced DNA condensation, which connects the condensation free energy with the number of ions in the external binding shell of the DNA duplex. Use of the model, instead of the more standard potential of mean force approach, makes the analysis computationally feasible for long enough duplexes and a variety of conditions explored in this work. According to the calculations based on the model, the DNA aggregation free energy in the presence of Na^+^ is at least 0.2 *k_B_T* per base pair more favorable that in the presence of K^+^. As a result, the attraction between the duplexes caused by the sodium ions is higher than that of potassium ions. Thus, our simulations combined with the multi-shell model, provide a quantitative explanation for the experimentally characterized difference between concentrations of monovalent cation concentrations needed to condense the DNA in the presence of PEG.

Our computations have revealed a noticeable dependence of the number of ions in the intersection of external ion binding shells on DNA sequence. Namely, more sodium ions bind in the external shell of the polyA homopolymer duplex compared to a GC-rich duplex. For potassium, this sequence effect is not as significant as for sodium. Thus, we make an easily testable prediction that in a crowded (PEG) environment, polyA DNA condenses more easily than mixed sequence DNA in the presence of sodium ions, while in solution with potassium ions the sequence dependence nearly disappears. We have suggested a plausible mechansim for the sequence dependence.

Another set of testable predictions is related to the interplay between the crowding agent and the specific ion type used to condense the DNA. We find that, depending on the ion affinity for the croding agent, it may “sequester” enough ions from the solution to effectively lower their ability to condense DNA. We predict that the propensity of an ion to condense the DNA depends on the type of crowding agent used in experiment or simulation. High affinity of the condesning ion to the crowding agent may lead to a counterintuitive effect – the addition of the crowding agent may oppose DNA condesnation, rather than promoting it, as could be expected from the general “reduced volume” argument.

This work has multiple potential implications, and offers a number of possible extensions beyond its original goal of explaining a curious phenomenon. Packaging of nucleic acids in a crowded environment is key to biology. For example, the genetic material is very tightly packed inside the viral capsid. To infect the cell, the virus has to cross from a sodium-rich plasma to a potassium-rich intra-cellular environment, which may affect the packing.

Perhaps even more intriguing are possible implications/extension of this work to chromatin packing, which obviously occurs in a highly crowded environment. Note that the DNA strands in the nucleosome, as well as in various tightly packed multi-nucleosome structures, come within the short enough distance that counterion condensation effects become relevant. Specifically, the nucleosomal forms “supergrooves”;^80^ parallel DNA fragments in the nucleosome structure have similar configurations to the one examined in this work for pairs of duplexes. Therefore, the mechanism of ion-induced DNA condensation under crowded conditions may also facilitate the DNA packing into the nucleosome.

Fluctuations of Na^+^ concentration during the cell cycle were already demonstrated:^87^ peaks of the sodium concentration were observed during mitosis and DNA synthesis phases. According to our model, an increase in the sodium concentration might facilitate chromatin condensation, in sequence-dependent manner. One may speculate if this effect might facilitate cell division during mitosis.

Finally, we note that NaCl, instead of KCl, has often been used as the main component of buffers employed in experiments and simulations of chromatin components, including the nucleosome. This work provides yet another argument for why one should consider this “detail” carefully, in designing experiments and simulations.

## 4. Methods

### 4.1. All-atom MD simulations

#### Structure preparation and simulation set-up

All MD simulations were carried out using ff99bsc1 force field and AMBER 18. ^88^ The DNA sequences were constructed in canonical B-form using Nucleic Acid Builder (NAB).^89^ Unless otherwise specified, single poly(dA_25_)·poly(dT_25_) fragment was used for simulations with a single DNA duplex, which we call “polyA”. We used 2 poly(dA_25_)·poly(dT_25_) duplexes in every system with two DNA duplexes (unless otherwise specified). The DNA duplexes in pairs are oriented as in Ref.,^69^ their grooves forming supergrooves. This specific orientation^69^ was associated with DNA-DNA attraction. DNA in the nucleosome structure forms supergrooves, ^80^ so the orientation is also biologically relevant. Each pair of DNA duplexes, and each single DNA duplex, was solvated with 13000*±* 30 water molecules, and 360 PEG 194 molecules. This number of polymer molecules is equal to 230 g/L concentration of PEG. For simulations with 28 g/L PEG, the number of polymer molecules was 44, and we added 16510*±* 30 molecules of water. For simulations without PEG, DNA duplexes were solvated with 17000 *±* 30 molecules of water. PEG molecule was constructed using tleap and antechamber tools of AMBER.^88^ We used OPC water model along with Joung/Cheatham ion parameters^90^ (ion type, *R_min_/*2(Å), *E* (kcal/mol)) (Na^+^, 1.226, 0.1684375; K^+^, 1.590, 0.2794651; Cl*^−^*, 2.760, 0.0116615). This solvent model was found to be a reasonable compromise for simulations of DNA in the ionic solution. ^41^ To calculate how many ions should be added to the system to reach the desired bulk salt concentration, we used SLTCAP^91^ method. For systems with a single 25 bp DNA, that approach gave 178 Na^+^ or K^+^ and 130 Cl*^−^* to reach 0.5 M bulk salt concentration. For systems with two DNA fragments, the numbers of added ions were (concentration (cations, anions)): 0.15 M (114, 18); 0.5 M (207, 111); 1 M (356, 260); 2 M (657, 561).

#### Simulation protocols

First we ran 2000 steps-long initial minimization using the steepest descent algorithm for the first 1000 steps and the conjugate gradient algorithm for 1000-2000 steps. After initial minimization, all the systems were heated from 0 to 300 K in canonical ensemble (NVT) over 18 ps, and then equilibrated for 2 ps in the same ensemble using 2 fs time step. Periodic boundary conditions and the particle mesh Ewald method were used. Frames were made every 10 ps in each production trajectory.

#### Single DNA duplex

After heating, each system with single DNA duplex was equilibrated for 40.06 ns in the isothermal-isobaric ensemble (NPT), using the same time step and Langevin dynamics with the collision frequency of 2*^−^*^1^ ps, to reach 1 atm pressure. After the equilibration, 100 ns long production trajectories were generated for each system using NPT ensemble and 2 fs time step. During the heating, equilibration and production simulation, 13th pairs of nucleotides in the systems with single 25 bp DNA were restrained to their original positions with 1 kcal/mol/A force constant to prevent drift of the duplex. Results, produced by this protocol, are described in Sec. 2.1.

#### Pairs of DNA duplexe

After heating each system with pair of DNA duplexes was equilibrated for 60 ps in isothermal–isobaric ensemble (NPT). During energy minimization, heating and first 60 ps of equilibration pairs of DNA were restrained to initial structure with 50.0 kcal/mol/A force constants to prevent significant DNA movement. Then we have aligned the initial structure of pair of DNA to the pair displaced during equilibraion. During the next 24 ps of equilibration, pairs of DNA were restrained with 0.01 kcal/mol/A force constants to the alligned structure to restore the inter-duplex distance. During the next 40 ns of equlibration DNA was restrained with 2 kcal/mol/A, and during the 100 ns-long production simulations the restraing force was 0.0001 kcal/mol/A force constants to the aligned DNA structure. We restrained the DNA duplexes to the structure with the inter-duplex distance of 24 Å unless otherwise specified. The force constant of restraints used during the production phase of the simulation (0.0001 kcal/mol/A) was chosen to allow DNA to drift apart to reach the distance of about 28 Å between axes but not to move far away from each other. When inter-duplex distance reach *∼* 28 Å energy of restraints is *∼* 1*k_B_T*. Results, produced by simulations with this protocol, are described in Sec. 2.2.1 and Sec. 2.2.2. For the computational experiment carried out to demonstrate DNA condensation, inter-duplex distance in the structure, DNA duplexes were constrained to, is 28 Å. In this case energy of restraints reach *∼* 1*k_B_T* when duplexes approach each other to a distance of *∼* 24 Å which is expected distance in condensed state. Results, produced by simulations with this protocol, are described in Sec. 2.2.3. The overlapping ion binding shells in this configuration are presented in Figure 1a. DNA duplexes were weakly restrained to reference structure to allow attraction but prevent a change of mutual orientation.

### 4.2. Trajectory Analysis and Calculation of Ion Distributions

For analysis of ion distribution around DNA, we use Curves+ and Canion.^92^ Curves+ translates coordinates into a convenient curvilinear helicoidal system. Canion analyzes the curvilinear coordinates of ions and generates density maps of ions around DNA. The output of Curves+ is also analyzed using package NumPy^93^ with Python 3. To compare the numbers of ions in the intersection of ion binding shells, we calculate the numbers of ions no further than 16 Å from both DNA helical axes; see below how we obtain the mean and the error of the computed values. To calculate the energy of electrostatic repulsion we used cpptraj plugin in AMBER, we do not take into account any molecules in the system except DNA in this calculations. To calculate the radial distribution function (RDF) of PEG to ions we used cpptraj program.

### 4.3. Multi-shell model of ion-induced nucleic acid condensation

The multi-shell model of ion-induced nucleic acid (NA) condensation was originally proposed^64^ to explain the difference between counterion-induced condensation propensities of dsRNA and dsDNA; the model was later expanded to provide quantitative expressions for the aggregation free energy of NA duplexes.^68^ Within the model, the space around NA’s helical axis is divided into three concentric ion binding shells: the area of deeply bound ions (0-7 Å from the helical axis), the internal shell (7-12 Å), and the external shell (12-16 Å). It was shown that the greater the number of ions in the external ion binding shell of the duplex, the more readily the condenses occurs.^64^ The ion-induced aggregation free energy can be represented as a sum of three additive components:^68^

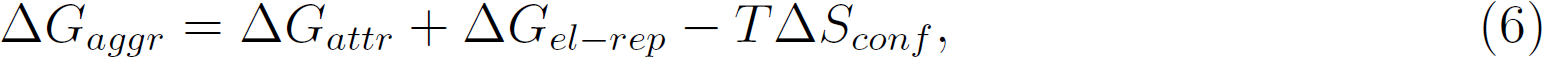

where Δ*G_attr_* is a short-range net attractive term, associated with the interaction of ions with DNA duplexes, Δ*G_el−rep_* represents the repulsion between duplexes, and the last term, *T* Δ*S_conf_* describes the loss of duplex configurational entropy (translational and rotational) because of the aggregation.^68^

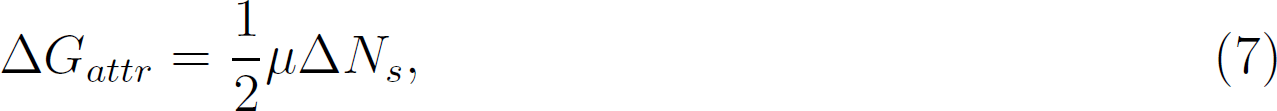

where *µ* is the binding affinity of ions to the intersection of the ion binding shells of adjacent DNA duplexes, and Δ*N_s_* is a number of ions in the overlapping shell region.

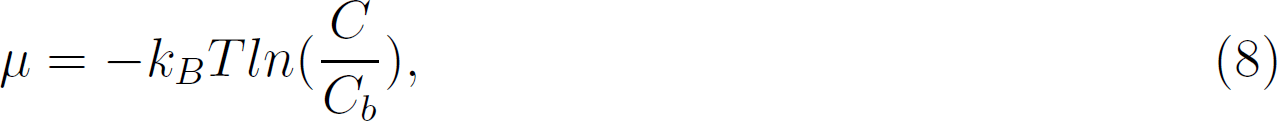

where *C* is the concentration of ions in the overlapping region of the ion binding shells, and *C_b_* is the bulk concentration of cations. In this work we calculate Δ*G_attr_* caused by the accumulation of ions in the intersection of external and internalion binding shells, see Figure 1b. The bulk concentration of ions was set using the SLTCAP method.^91^ To calculate the number of ions in the overlap region of the ion binding shells we count the number of ions that are not further than 16 Å from both helical axes of the DNA duplexes. To calculate the ion concentration, we divide the number of ions by the volume of the intersection. To estimate the volume, we calculate the inter-duplex distance (*d*), averaged over time and over the region of 3-23 nucleotides. Two nucleotide pairs were excluded from consideration from every end of DNA to avoid edge effects. Then we calculated the volume of area limited by surfaces of two parallel cylinders of radii 16 Å, length 82.5 Å(equal to the length of 25 bp-long DNA) and with the distance between axes equal to *d*.

### 4.4. Estimation of computed values and their statistical errors

For each calculated value for which we seek an estimate of its statistical error, the error is estimated with the slicing method.^94^ First, we count the number of ions in the intersection of ion binding shells in every snapshot along the trajectory. Then, we plot the distribution of the number of snapshots containing the given number of deeply bound ions. We find where the distribution reaches half of its maximum value; we take those two numbers of deeply bound ions as the thresholds, Figure 8. Then we count the number of pieces of the trajectory (*N*), in each of which the number of ions in the intersection of ion binding shells crosses each of the threshold values at least once. After that, we cut the trajectory onto *N* equal fragments, and calculate the corresponding average number of ions and other reported values. The standard deviation *σ* is computed, and the assigned error is the standard error of the mean 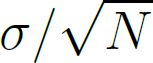.

**Figure 8:**
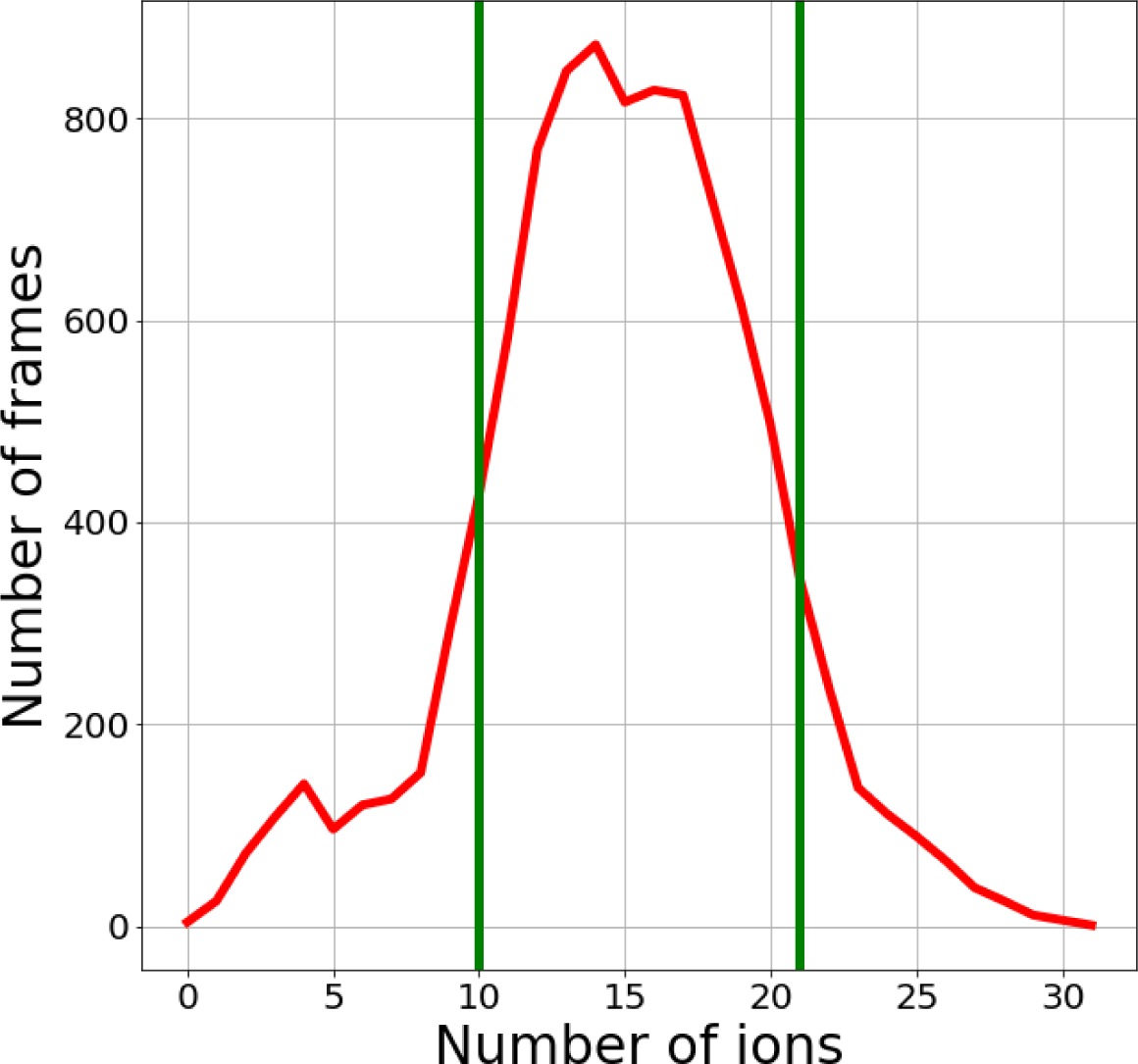
Shown is a dependence of the number of snapshots containing the given number of ions in the intersection of ion binding shells. Taken from the trajectory of polyA DNA duplexes with reference inter-DNA distance of 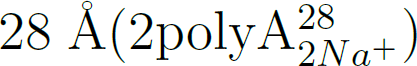. Vertical green lines intersect the distribution curve at half its maximum value.

## Acknowledgement

Y.S.K. and I.Yu.G. were supported by the Ministry of Science and Higher Education of the Russian Federation (agreement 075-03-2023-106, project FSMG-2021-0002). P. Z. acknowledges support from the Russian Science Foundation (project number: 21-79-20228).

## Supporting Information Available

Supporting Text, Figures S1-S2 and Tables S1-S8 are afialable at…

## Graphical TOC Entry

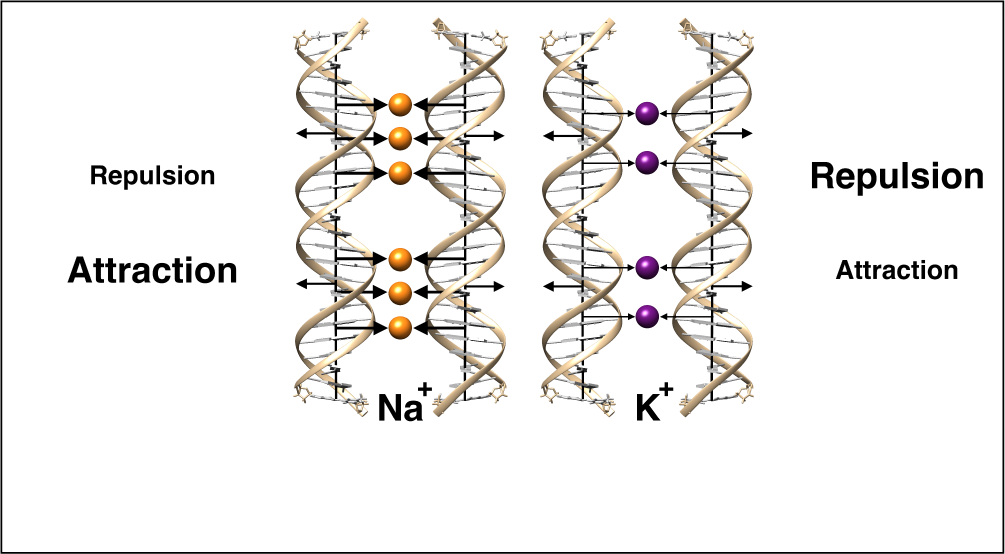

