## Supplemental Info for "Why Na+ has higher propensity than K+ to condense DNA in a crowded environment"

### Supplementary information for: Why $Na^+$ has higher propensity than $K^+$ to condense DNA in a crowded environment

Egor S. Kolesnikov,[†] Ivan Yu. Gushchin,[†] Petr A. Zhilyaev,[‡] and Alexey V. Onufriev[*,¶,§,‖]

†*Research Center for Molecular Mechanisms of Aging and Age-Related Diseases, Moscow Institute of Physics and Technology, Dolgoprudny 141700, Russia*

‡*Center for Design, Manufacturing and Materials, Skolkovo Institute of Science and Technology, Bolshoy Boulevard 30, bld. 1, Moscow 121205, Russia*

¶*Department of Computer Science, Virginia Tech, Blacksburg 24061-0131, United States*

§*Department of Physics, Virginia Tech, Blacksburg 24061-0131, United States*

‖*Center for Soft Matter and Biological Physics, Virginia Tech, Blacksburg 24061-0131, United States*

Supporting Text

Supporting Figures S1-S2

Supporting Tables S1-S8

In Figures S1 and S2 we examine, in some detail, the differences between how sodium and potassium interact with PEG. In the absence of PEG, the numbers of $Na^+$ and $K^+$ in the external ion binding shell differ less ($25.3 \pm 0.2$ $Na^+$ vs. $23.3 \pm 0.2$ $K^+$) than they do in solutions with 230 g/L PEG ($28.4 \pm 0.2$ $Na^+$ vs. $22.3 \pm 0.2$ $K^+$), see Main Text Figure 7. The similarity of the numbers of $Na^+$ and $K^+$ in the external shell in PEG-free solution means that the ion affinity to the external ion binding shell of DNA does not differ significantly between $Na^+$ and $K^+$. Comparison of ion and PEG distributions in two types of solvent reveals the origins of this difference in the behavior of these two ions in the presence and absence of PEG. One can see in Figure S2 that after the addition of PEG, the peaks of ion distribution do not change their locations, but the density of PEG around $K^+$ ions is higher than around $Na^+$, see Figure S3. Therefore, $K^+$ chemical potential in solution with PEG is less than that of $Na^+$. The difference between $Na^+$ and $K^+$ chemical potentials in solution with PEG causes the difference in the numbers of sodium and potassium ions in the external ion binding shell of the DNA in the presence of PEG. Fewer potassium ions than sodium ions are accumulated in the external ion binding shell of single DNA in the presence of PEG.

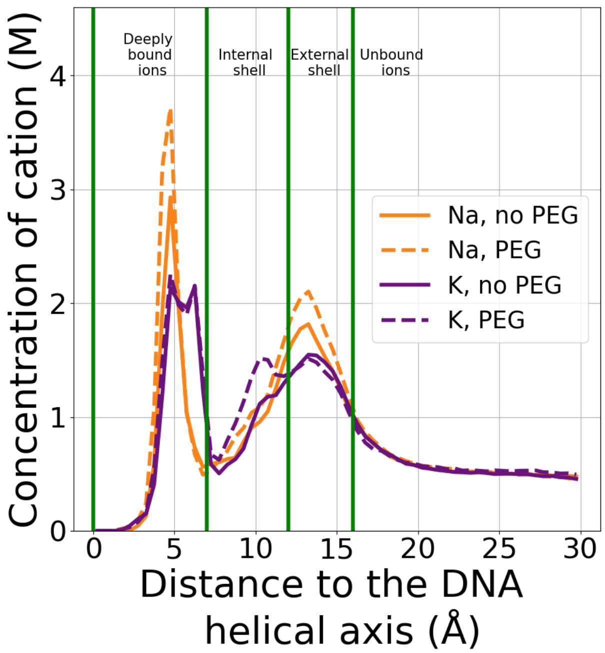


Figure S1: Locations of the peaks of ion distributions around DNA do not depend on the presence of PEG. Shown are distributions of $Na^+$ (orange) and $K^+$ (purple) ions around single DNA duplex in the presence (dashed) and absence (solid) of PEG around polyA DNA. Bulk concentration of NaCl and KCl is 0.5M.

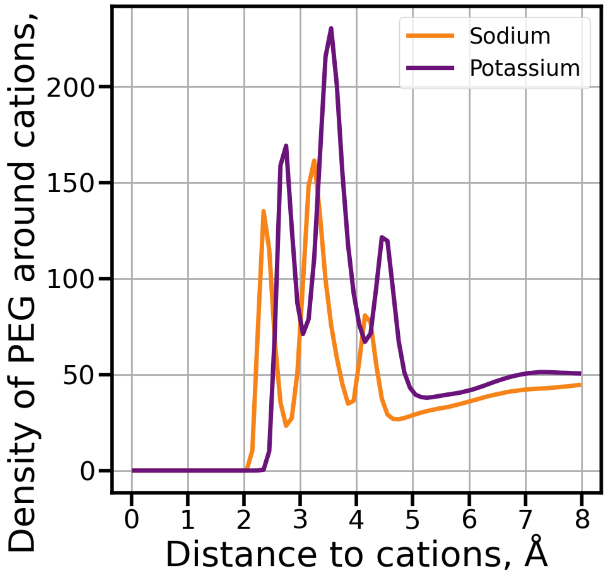


Figure S2: Density of PEG around potassium ions is higher than that for sodium ions. This difference points to higher affinity of PEG to $K^+$ than to $Na^+$. Shown are radial distribution functions of PEG around sodium ions (orange) and potassium ions (purple). Bulk concentration of NaCl and KCl is 0.5M, bulk PEG concentration is 230 g/L.

Table S1: Identifiers of all the simulations carried out in this work.

| Number of duplexes | DNA sequence | Reference distance, Å | Cation type | PEG concentration, g/L | Salt concentration, M | | | |
|---|---|---|---|---|---|---|---|---|
| | | | | | 0.15 | 0.5 | 1 | 2 |
| 1 | polyA | - | $Na^+$ | 0 | | (1polyA$_{0.5Na^+}$; 0 PEG) | | |
| | | | | 28 | | (1polyA$_{0.5Na^+}$; 28 PEG) | | |
| | | | | 230 | | 1polyA$_{0.5Na^+}$ | | |
| | | | $K^+$ | 0 | | (1polyA$_{0.5K^+}$; 0 PEG) | | |
| | | | | 28 | | (1polyA$_{0.5K^+}$; 28 PEG) | | |
| | | | | 230 | | 1polyA$_{0.5K^+}$ | | |
| | | | $Rb^+$ | 230 | | 1polyA$_{0.5Rb^+}$ | | |
| | mixed | | $Na^+$ | | | 1mixed$_{0.5Na^+}$ | | |
| | | | $K^+$ | | | 1mixed$_{0.5K^+}$ | | |
| 2 | polyA | 24 | $Na^+$ | | 2polyA$^{24}_{0.15Na^+}$ | 2polyA$^{24}_{0.5Na^+}$ | 2polyA$^{24}_{1Na^+}$ | 2polyA$^{24}_{2Na^+}$ |
| | | | $K^+$ | | 2polyA$^{24}_{0.15K^+}$ | 2polyA$^{24}_{0.5K^+}$ | 2polyA$^{24}_{1K^+}$ | 2polyA$^{24}_{2K^+}$ |
| | | 28 | $Na^+$ | | | | | 2polyA$^{28}_{2Na^+}$ |
| | | | $K^+$ | | | | | 2polyA$^{28}_{2K^+}$ |
| | mixed | 24 | $Na^+$ | | | | | 2mixed$^{24}_{2Na^+}$ |
| | | | $K^+$ | | | | | 2mixed$^{24}_{2K^+}$ |
| | | 28 | $Na^+$ | | | | | 2mixed$^{28}_{2Na^+}$ |
| | | | $K^+$ | | | | | 2mixed$^{28}_{2K^+}$ |

Table S2: Number of sodium ions in the intersection of ion binding shells is higher than that of potassium in a wide range of salt concentrations. Listed are numbers of sodium and potassium ions in the intersection of ion binding shells belonging to two DNA duplexes. Reference inter-duplex distance 24 Å.

| | Salt concentration, M | Number of ions | | Difference of number of ions |
|---|---|---|---|---|
| | | Na | K | |
| All ion binding shells | 0.15 | 7.9 ± 0.2 | 4.9 ± 0.2 | 2.9 ± 0.3 |
| | 0.5 | 10.3 ± 0.5 | 7.8 ± 0.4 | 2.5 ± 0.6 |
| | 1 | 13.2 ± 0.4 | 10.2 ± 0.3 | 3.0 ± 0.5 |
| | 2 | 18.0 ± 0.3 | 12.6 ± 0.4 | 5.4 ± 0.5 |
| internal and external ion binding shells | 0.15 | 1.8 ± 0.2 | 1.0 ± 0.1 | 0.8 ± 0.2 |
| | 0.5 | 3.0 ± 0.4 | 2.3 ± 0.6 | 0.7 ± 0.6 |
| | 1 | 4.2 ± 0.4 | 3.6 ± 0.5 | 0.6 ± 0.5 |
| | 2 | 7.9 ± 0.4 | 4.2 ± 0.4 | 3.7 ± 0.6 |

Table S3: Energy of electrostatic repulsion between two DNA duplexes does not depend on the type of cation significantly. Listed are repulsion energies of DNA duplexes in different conditions. Ions between DNA duplexes are not taken into consideration. Reference inter-duplex distance 24 Å.

| Salt concentration, M | $\Delta G_{el-rep}$, $k_B$T | | $\Delta\Delta G_{el-rep}$ |
|---|---|---|---|
| | Na | K | |
| 0.15 | 22.76 ± 0.03 | 22.51 ± 0.03 | 0.26 ± 0.04 |
| 0.5 | 22.99 ± 0.05 | 22.81 ± 0.05 | 0.18 ± 0.07 |
| 1 | 23.00 ± 0.04 | 23.07 ± 0.04 | -0.07 ± 0.05 |
| 2 | 23.35 ± 0.03 | 22.98 ± 0.04 | 0.36 ± 0.05 |

Table S4: Attraction of duplexes is more energetically beneficial in solution with sodium than in the presence of potassium in a wide range of salt concentrations. Listed are energies of attraction between two DNA duplexes caused by ions in the intersection of external ion binding shells. Calculated using multi-shell model of ion-induced DNA condensation.

| Salt concentration, M | $\Delta G_{attr}$, $k_B$T | | $\Delta\Delta G_{attr}$ |
|---|---|---|---|
| | Na | K | |
| 0.15 | -22.4 ± 0.6 | -12.9 ± 0.5 | -9.6 ± 0.8 |
| 0.5 | -18.0 ± 0.8 | -12.1 ± 0.6 | -6 ± 1 |
| 1 | -16.6 ± 0.5 | -9.5 ± 0.3 | -7.1 ± 0.6 |
| 2 | -11.5 ± 0.2 | -6.2 ± 0.2 | -5.3 ± 0.3 |

#### Convergence of the computed values

We varied the number of water and PEG molecules in the simulations with 2 DNA duplexes 4-fold, and the simulation time 10-fold, to check if the computed numbers of ions and other parameters change. For this test, we chose the systems with polyA DNA and 1M bulk salt concentration. We quadrupled number of water molecules and PEG by nearly doubling the dimensions of the box in the directions perpendicular to the helical axis of the DNA. These systems required extra 270 ps of equilibration using CPU. For the system in the standard box we extended the simulation time to 1 microsecond, and compared the number of ions in the intersections of ion binding shells and energies obtained from the first 100 ns of the trajectory with those averaged over the whole trajectory. Comparison of the calculated values averaged over different time windows, and with different sizes of the solvent box, are presented below.

Table S5: Numbers of sodium and potassium ions in the intersection of ion binding shells belonging to two DNA duplexes and energies in systems with different numbers of solvent molecules do not differ significantly. Bulk NaCl and KCl concentration is 1 M.

| Value | Number of solvent molecules | Ion type | | Difference of value |
|---|---|---|---|---|
| | | $Na^+$ | $K^+$ | |
| Number of ions | standard | 13.2 ± 0.4 | 10.2 ± 0.3 | 3.0 ± 0.5 |
| | quadrupled | 13.6 ± 0.4 | 10.4 ± 0.3 | 3.2 ± 0.5 |
| $\Delta G_{attr}$, $k_B$T | standard | -16.6 ± 0.5 | -9.5 ± 0.3 | -7.1 ± 0.6 |
| | quadrupled | -15.4 ± 0.4 | -10.1 ± 0.3 | -5.2 ± 0.5 |
| $\Delta G_{el-rep}$, $k_B$T | standard | 23.00 ± 0.04 | 23.07 ± 0.04 | -0.07 ± 0.05 |
| | quadrupled | 23.23 ± 0.03 | 22.96 ± 0.04 | 0.27 ± 0.05 |

Table S6: Numbers of sodium and potassium ions in the intersection of ion binding shells belonging to two DNA duplexes and energies in 100 and 1000 ns trajectories do not differ significantly. Bulk NaCl and KCl concentration is 1 M.

| Value | Simulation time, ns | Ion type | | Difference of value |
|---|---|---|---|---|
| | | $Na^+$ | $K^+$ | |
| Number of ions | 100 | 13.2 ± 0.4 | 10.2 ± 0.3 | 3.0 ± 0.5 |
| | 1000 | 13.0 ± 0.1 | 10.2 ± 0.1 | 2.8 ± 0.2 |
| $\Delta G_{attr}$, $k_B T$ | 100 | -16.6 ± 0.5 | -9.5 ± 0.3 | -7.1 ± 0.6 |
| | 1000 | -15.4 ± 0.2 | -10.7 ± 0.2 | -4.8 ± 0.3 |
| $\Delta G_{el-rep}$, $k_B T$ | 100 | 23.00 ± 0.04 | 23.07 ± 0.04 | -0.07 ± 0.05 |
| | 1000 | 23.12 ± 0.02 | 23.00 ± 0.02 | 0.15 ± 0.03 |

### Dependence of the computed values on the DNA sequence

Table S7: Number of sodium ions and attraction energies depend on the DNA sequence noticeably. Listed are number of ions in the intersection of ion binding shells, energy of repulsion and attraction and average inter-duplex distances in the systems of adjacent polyA duplexes and mixed-sequence duplexes. Reference distance is 24 Å. Bulk NaCl and KCl concentration is 2 M.

| sequence | Ion type | Number of ions in the intersection of ion binding shells | $\Delta G_{el-na}$, $k_B T$ | $\Delta G_{attr}$, $k_B T$ | Inter-duplex distance, Å |
|---|---|---|---|---|---|
| polyA | $Na^+$ | 18.0 ± 0.3 | 23.35 ± 0.03 | -11.5 ± 0.2 | 25.36 ± 0.25 |
| | $K^+$ | 12.6 ± 0.4 | 22.98 ± 0.04 | -6.2 ± 0.2 | 26.4 ± 0.2 |
| | difference | 5.4 ± 0.5 | 0.36 ± 0.05 | -5.3 ± 0.7 | -1.04 ± 0.32 |
| mixed | $Na^+$ | 13.4 ± 1.5 | 25.6 ± 0.1 | -7.5 ± 0.8 | 26.37 ± 0.25 |
| | $K^+$ | 11.5 ± 0.7 | 25.5 ± 0.1 | -7.3 ± 0.5 | 26.96 ± 0.25 |
| | difference | 1.9 ± 1.6 | -0.05 ± 0.1 | -0.2 ± 0.9 | -0.59 ± 0.35 |

Table S8: Number of sodium ions and attraction energies depend on the DNA sequence noticeably. Listed are number of ions in the intersection of ion binding shells, energy of repulsion and attraction and average inter-duplex distances in the systems of adjacent polyA duplexes and mixed-sequence duplexes. Reference distance is 28 Å. Bulk NaCl and KCl concentration is 2 M.

| sequence | Ion type | Number of ions in the intersection of ion binding shells | $\Delta G_{el-rep}$, $k_BT$ | $\Delta G_{attr}$, $k_BT$ | Inter-duplex distance, Å |
|---|---|---|---|---|---|
| polyA | $Na^+$ | 15 ± 1 | 22.98 ± 0.08 | -10.0 ± 0.7 | 26.4 ± 0.4 |
| | $K^+$ | 8.2 ± 0.5 | 22.49 ± 0.06 | -4.7 ± 0.3 | 28.16 ± 0.23 |
| | difference | 6.7 ± 1.1 | 0.5 ± 0.1 | -5.2 ± 0.7 | -1.68 ± 0.46 |
| mixed | $Na^+$ | 11.6 ± 0.3 | 25.41 ± 0.04 | -7.7 ± 0.2 | 27.06 ± 0.25 |
| | $K^+$ | 8.7 ± 0.4 | 25.27 ± 0.05 | -4.5 ± 0.2 | 27.62 ± 0.25 |
| | difference | 2.9 ± 0.5 | 0.13 ± 0.06 | -3.2 ± 0.3 | -0.56 ± 0.35 |

We have also re-examined the case of a single DNA duplex described in the first part of the Results section of the main text. The following simulations have been performed: 1mixed$_{0.5Na^+}$, 1mixed$_{0.5K^+}$. In the simulations with a single DNA duplex, the number of sodium ions in the external shell of DNA is higher than that of potassium ($Na^+$: $26.0 \pm 0.3$, $K^+$: $20.1 \pm 0.2$), and the estimated $\Delta\Delta G_{attr}(Na^+ \ vs. \ K^+) = 0.15 k_BT$ per base pair for hexagonal packing of DNA, compared with $0.2 k_BT$ for polyA, see main text. Thus, the single duplex approach is sensitive enough to correctly predict that sodium is more potent than potassium at condensing DNA in the case of the mixed sequence (we have seen it already for polyA DNA, see main text).

However, a more detailed analysis shows that not all of the conclusions made using the more accurate two-duplex systems can be reached via more efficient single duplex simulations. Namely, one can see that the numbers of both sodium and potassium in the external shell of a single mixed-sequence DNA duplex is less than the numbers of ions in the external shell of polyA DNA (Na: $26.0 \pm 0.3$ for mixed sequence vs. $28.4 \pm 0.2$ for polyA, K: $20.1 \pm 0.2$ for mixed sequence vs. $22.3 \pm 0.2$ for polyA). Note that for pairs of duplexes, the same statistically significant trend is observed only for $Na^+$, but not for $K^+$, see main text. This difference indicates that the distributions of ions change in some (albeit subtle) way when two DNA duplexes approach each other. That is as far as monovalent ions are concerned, single duplex simulations do not always provide accurate enough picture of the counterion distributions.
